# Early-Stage Ocular Hypertension Alters Retinal Ganglion Cell Synaptic Transmission in the Visual Thalamus

**DOI:** 10.1101/669135

**Authors:** Ashish Bhandari, Jennie C. Smith, Yang Zhang, Lisa Reid, Toni Goeser, Shan Fan, Deepta Ghate, Matthew J. Van Hook

## Abstract

Axonopathy is a hallmark of many neurodegenerative diseases including glaucoma, where elevated intraocular pressure (ocular hypertension, OHT) stresses retinal ganglion cell (RGC) axons as they exit the eye and form the optic nerve. OHT causes early changes in the optic nerve such as axon atrophy, transport inhibition, and gliosis. Importantly, many of these changes appear to occur prior to irreversible neuronal loss, making them promising points for early diagnosis of glaucoma. It is unknown whether OHT has similarly early effects on the function of RGC output to the brain. To test this possibility, we elevated eye pressure in mice by anterior chamber injection of polystyrene microbeads. 5 weeks post-injection, bead-injected eyes showed a modest RGC loss in the peripheral retina, as evidenced by RBPMS antibody staining. Additionally, we observed reduced dendritic complexity and lower spontaneous spike rate of On-αRGCs, targeted for patch clamp recording and dye filling using a *Opn4*-cre reporter mouse line. To determine the influence of OHT on retinal projections to the brain, we expressed Channelrhodopsin-2 (ChR2) in melanopsin-expressing retinal ganglion cells by crossing the *Opn4*-cre mouse line with a ChR2-reporter mouse line and recorded post-synaptic responses in thalamocortical relay neurons in the dorsal lateral geniculate nucleus (dLGN) of the thalamus evoked by stimulation with 460 nm light. The use of a *Opn4*-cre reporter system allowed for expression of ChR2 in a narrow subset of RGCs responsible for image-forming vision in mice. Five weeks following OHT induction, paired pulse and high-frequency stimulus train experiments revealed that presynaptic vesicle release probability at retinogeniculate synapses was elevated. Additionally, miniature synaptic current frequency was slightly reduced in brain slices from OHT mice and proximal dendrites of post-synaptic dLGN relay neurons, assessed using a Sholl analysis, showed a reduced complexity. Strikingly, these changes occurred prior to major loss of RGCs labeled with the *Opn4*-Cre mouse, as indicated by immunofluorescence staining of ChR2-expressing retinal neurons. Thus, OHT leads to pre- and post-synaptic functional and structural changes at retinogeniculate synapses. Along with RGC dendritic remodeling and optic nerve transport changes, these retinogeniculate synaptic changes are among the earliest signs of glaucoma.

## Introduction

Axonopathy is a common feature of many neurodegenerative diseases including glaucoma, a blinding disorder that leads to vision loss due to the degeneration of retinal ganglion cells (RGCs) and their projections to visual nuclei in the brain (Calkins, 2012; Weinreb et al., 2014). Intraocular pressure (IOP) is the only controllable risk factor for glaucoma and high IOP (ocular hypertension, OHT) triggers a cascade of pathological events by stressing retinal ganglion cell (RGC) axons as they converge at the optic nerve head to form the optic nerve. The development of tools for diagnosing and treating glaucoma prior to irreversible RGC loss hinges on an understanding of how the disease processes traumatize the optic nerve and influence the performance of neurons throughout the visual pathway.

OHT leads to changes in the structure and function of retinal neurons prior to somatic degeneration such as dendritic pruning, changes in excitability, and synaptic dysfunction (Della Santina et al., 2013; El-Danaf and Huberman, 2015; Frankfort et al., 2013; Ou et al., 2016; Pang et al., 2015; Park et al., 2014; Risner et al., 2018). In the optic nerve, OHT disrupts retrograde and anterograde transport along RGC axons in both inducible and inherited rodent glaucoma models and these transport changes appear to contribute to eventual neuronal loss (Pease et al., 2000; Quigley et al., 2000; Crish et al., 2010, 2013; Dengler-Crish et al., 2014; Smith et al., 2016). In retinorecipient brain regions such as the superior colliculus (SC), suprachiasmatic nucleus (SCN), and dorsal lateral geniculate nucleus of the thalamus (dLGN), studies with post-mortem human tissue and animal models have shown that OHT and glaucoma eventually lead to glial activation, changes in neuronal structure, neuron loss, and atrophy of RGC axon terminals (Calkins, 2012; Chaturvedi et al., 1993; Chen et al., 2015; Crish et al., 2010; Gupta et al., 2006, 2007, 2009; Ito et al., 2011; Liu et al., 2014; Ly et al., 2011; Shimazawa et al., 2012; Smith et al., 2016; Wong and Brown, 2012; Yücel et al., 2000, 2001, 2003, 2006; Yucel and Gupta, 2015; Zhang et al., 2009).

OHT-dependent changes in optic nerve function are likely to trigger homeostatic and/or pathological changes in regions receiving retinal inputs. Extracellular recordings in the mouse SC have shown that OHT alters receptive fields of SC neurons receiving retinal signals (Chen et al., 2015). Still, it is largely unknown how OHT influences RGC synaptic outputs in the brain. The goal of this study was to determine how early-stage and modest elevation of eye pressure of the sort that mimics human disease (Mao et al., 1991) affects excitatory synaptic output from RGCs in the dLGN.

Using an optogenetic mouse reporter to drive Channelrhodopsin-2 (ChR2) in a small subset of RGCs in tandem with brain slice electrophysiology, we determined that OHT increased the probability of action potential-triggered synaptic vesicle exocytosis from RGC axon terminals and caused detectable changes in the frequency of miniature synaptic events and dendritic structure of neurons in the dLGN. These changes occurred prior to major RGCs loss, as indicated by immunofluorescent staining of RGC somata. These novel findings indicate that changes to the performance of RGC synapses with TC neurons in the dLGN, along with RGC dendritic remodeling and axon transport deficits, are among some of the earliest functional changes in the visual pathway accompanying elevation of eye pressure in glaucoma.

## Materials and Methods

### Animals and microbead occlusion model

Animal protocols were approved by the Institutional Animal Care and Use Committee at the University of Nebraska Medical Center. To explore a subset of retinal neurons signaling to the dLGN, we made use of a mouse line in which Cre recombinase is knocked into the alleles for melanopsin (*Opn4*). While M1-type melanopsin-expressing RGCs are well known for their role in circadian photoentrainment and pupillary constriction, these cells generally do not project to the dLGN. Several other melanopsin-expressing RGCs including M4, M5, and M6 RGCs have low levels of melanopsin expression, receive substantial rod- and cone-driven synaptic inputs, project to the dLGN, and are involved in conscious vision (Ecker et al., 2010; Estevez et al., 2012; Quattrochi et al., 2019; Schmidt et al., 2014; Stabio et al., 2018). For instance, the M4-type melanopsin RGC is identical with the on-sustained αRGC (Estevez et al., 2012; Schmidt et al., 2014). Thus, we opted to use the *Opn4*-Cre mouse line simply as a convenient way to drive reporter expression in a subpopulation of dLGN-projecting RGCs.

Channelrhodopsin-2 (ChR2) was expressed in melanopsin-expressing retinal neurons by crossing the *Opn4*-Cre/Cre mouse (Ecker et al., 2010) with the Ai32 reporter line (Jackson Labs # 024109) (Madisen et al., 2012). In these *Opn4*-Cre/+;Ai32 mice, Cre-mediated excision of a LoxP-flanked stop codon leads to expression of a ChR2-enhanced yellow fluorescent protein fusion protein (ChR2-EYFP) under control of a CAG promoter. *Opn4*-Cre/+;Ai32 mice were used for brain slice electrophysiology experiments and retinal immunofluorescence staining for performing RGC density counts. For patch-clamp recordings of RGC light responses and measurements of RGC dendritic complexity, we used *Opn4*-Cre/Cre mice crossed with the Z/EG reporter line (Novak et al., 2000). In those *Opn4*-cre/+;Z/EG mice, enhanced green fluorescent protein (EGFP) is expressed in melanopsin-expressing RGCs. C57Bl/6J mice (Jackson Labs #00664) were used for immunofluorescence staining in dLGN brain slices.

To induce ocular hypertension, we performed bilateral injections of fluorescently-tagged polystyrene microspheres (10 micron, Invitrogen F8836) into the anterior chambers of 6-8 weeks old mice (Calkins et al., 2018; Sappington et al., 2010). To accomplish this, mice were anesthetized with isoflurane and topical anesthetic (0.5% proparacaine, Akorn, Lake Forest, IL) and dilating (1% tropicamide, Akorn) eye drops were instilled into each eye. A small volume of beads (~1-2 μL) that had been concentrated to approx. 14×10^6^ beads/mL by microcentrifuging and resuspending were injected into the eye via a tapered glass pipette with an opening of ~50-100 microns connected to a manual microsyringe pump via mineral oil-filled tubing. Control mice received bilateral anterior chamber injections with sterile saline. Following injection, each eye was instilled with antibiotic eye drops (0.3% ciprofloxacin, Sandoz, Princeton, NJ) and mice were allowed to recover from anesthesia on a warming pad before being returned to their home cage.

Intraocular pressure (IOP) measurements were performed before bead/saline injections and at 2 days, 1 week, 2 weeks, and 4 weeks post-injection. To do this, mice were lightly anesthetized with isoflurane and IOP was measured with a Tonolab rebound tonometer (iCare, Vantaa, Finland). The Tonolab tonometer takes a series of six measurements and reports the mean of the middle four. We took an average of three such measurements for each eye. Following IOP measurement, we administered hydrating eye drops to each eye (Systane, Alcon, Fort Worth, TX) as mice were allowed to recover from anesthesia.

### Optical coherence tomography (OCT) imaging

12 radial sections of the optic nerve were acquired from anesthetized mice with a Heidelberg Spectralis OCT (Heidelberg Engineering, Inc., Franklin, MA) adapted with a +35 diopter lens. Measurements of RNFL+GCL+IPL thickness using three sections over 360 degrees were performed by two graders blinded to treatment conditions at 500 microns from Bergmeister’s papillae-ILM junctions. If discrepancy between graders was >20 μm, a third grader measured the section. If all graders’ measurements were different, the section was considered ungradable. The grading protocol was as stringent as that used in human studies.

### Brain slice preparation

We prepared acute coronal brain slices for *in vitro* patch clamp recordings following a “protected recovery” method (Ting et al., 2014, 2018). Mice were euthanized by CO_2_ asphyxiation and cervical dislocation. They were then decapitated and brains were rapidly dissected into a slush of artificial cerebrospinal fluid (aCSF) containing (in mM) 128 NaCl, 2.5 KCl, 1.25 NaH_2_PO_4_, 24 NaHCO_3_, 12.5 glucose, 2 CaCl_2_, 2 MgSO_4_ and continuously bubbled with a mixture of 5% CO_2_ and 95% O_2_. The cerebellum was removed with a razor blade and the brain was affixed to the cutting chamber using cyanoacrylate glue. 250 micron thick coronal brain sections through the dLGN were cut with a vibrating microtome (Leica VT1000S) and hemisected through the midline before being transferred to a net submerged in a NMDG-based solution composed of (in mM) 92 NMDG, 2.5 KCl, 1.25 NaH_2_PO_4_, 25 glucose, 30 NaHCO_3_, 20 HEPES, 0.5 CaCl_2_, 10 MgSO_4_, 2 thiourea, 5 L-ascorbic acid, and 3 Na-pyruvate, warmed to ~30°C and bubbled with CO_2_/O_2_. After a 10 minute incubation in the NMDG solution, slices were transferred to a chamber containing room temperature aCSF and allowed to recover for one hour before beginning patch clamp experiments. 250 μm Parasagittal brain slices were prepared using the same solutions as for coronal slices following the cutting procedure described by Turner & Salt (1998). Briefly, the brain was hemisected with a razor blade angled ~5 degrees from the medial longitudinal fissure and ~20 degrees from the horizontal plane. The medial surface was affixed to the vibratome cutting stage to prepare parasagittal slices containing the dLGN and optic tract. Reagents were purchased from Thermo Fisher Scientific (Waltham, MA) unless noted otherwise.

### Electrophysiology

dLGN slices were positioned in a recording chamber on an upright fixed-stage microscope (Olympus BX51WI) and superfused by a gravity-fed system with room temperature aCSF (23-24°C) at 2-4 mL/min. The aCSF was supplemented with 60 μM picrotoxin. Thalamocortical (TC) relay neurons were targeted for whole-cell patch clamp recording in the dLGN core using patch pipettes pulled from thin-walled borosilicate glass capillary tubing with an internal filament. We used a Multiclamp 700A amplifier, a DigiData 1550B digitizer, and pClamp 10 electrophysiology software (Axon/Molecular Devices, San Jose, CA). Patch pipettes were positioned using Sutter MP-225 manipulators (Sutter Instruments, Novato, CA). The pipette solution contained (in mM) 120 Cs-methanesulfonate, 2 EGTA, 10 HEPES, 8 TEA-Cl, 5 ATP-Mg, 0.5 GTP-Na_2_, 5 phosphocreatine-Na_2_, 2 QX-314 (for voltage-clamp measurements of vesicle release probability). Reported voltages are corrected for a measured liquid junction potential 10 mV. Optogenetic stimulation of ChR2-expressing RGC axons was accomplished with a 460 nm full field flash generated by a TLED system (Sutter Instruments) and delivered through the microscope objective. The LED was triggered using a digital TTL pulse while LED intensity (0.05-1.5 mW) was adjusted by supplying an analog voltage output from the Digidata 1550B. TLED power was calibrated using a Metrologic digital laser power meter. In parasagittal sections, the optic tract was stimulated with 0.2-0.5 ms current pulses (1-500 μA) delivered from a bipolar stimulating electrode positioned in the optic tract ventral to the ventral lateral geniculate nucleus (Chen and Regehr, 2000).

During recording, the aCSF was supplemented with 100 μM cyclothiazide (Santa Cruz, Dallas, TX) and 200 μM γ—D-glutamylglycine (γDGG, Abcam, Cambridge, MA) for release probability and mEPSC measurements. For strontium substitution experiments, we replaced 2 mM CaCl_2_ in the ACSF with 3 mM SrCl_2_. Other pharmacological agents (i.e. CNQX, TTX, picrotoxin) were diluted from 500-2000x stock solutions into ACSF and bath applied. CNQX and TTX were purchased from Tocris/Bio-Techne (Minneapolis, MN).

Miniature EPSCs (mEPSCs) were recorded using a 20-60 s duration recording in the absence of stimulation. For each cell, the first ~100 events were detected and analyzed using MiniAnalysis software (Synaptosoft Inc, Fort Lee, NJ).

For RGC recordings in retinal flat mounts, retinas were dissected free from the sclera in room temperature Ames’ medium bubbled with 5% CO_2_ in O_2_. Four relieving cuts were made in the peripheral retina, which was then positioned on a poly-D-lysine/laminin-coated coverslip (BD Biocoat) on a recording chamber, anchored down with a slice hold-down, and covered with an enzyme mixture (240 U/mL collagenase and 2 mg/mL hyaluronidase) diluted in Ames’ (Schmidt and Kofuji, 2011). The retina with enzyme mixture was incubated in a humidified darkened chamber saturated with 5%CO2/O2 at room temperature for 10 minutes before being positioned on the microscope stage and superfused with Ames at a rate of 2-4 mL/min. On sustained-αRGCs (OnαRGCs, a.k.a. M4-type melanopsin cells) (Estevez et al., 2012) were targeted based on somatic EGFP fluorescence in *Opn4*-cre/+;Z/EG mouse retinas, and large soma size (20+ microns diameter) and identity was confirmed following dye-filling. For M4 light responses and dendritic morphology experiments, we used the *Opn4*-cre/+;Z/EG mice and the pipette solution was Cs-based and was supplemented with Qx-314 (2 mM), CF568 dye (100 μM). Spontaneous spiking was measured in tight-seal cell attached mode prior to breaking in.

Clampfit 10 (Molecular Devices) and GraphPad Prism 7 (La Jolla, CA) were used for analysis of electrophysiology data.

### Dendritic imaging and Sholl analysis

For dendritic analysis, TC neurons in 250 micron brain slices or RGCs in flat-mount retinas were loaded by passive diffusion with 2% neurobiotin (Vector Labs, Burlingame, CA) and/or 100 μM CF568 dye (Biotium, Fremont, CA) dissolved in pipette solution during whole-cell recording. Following recording, brain slices were fixed in 4% paraformaldehyde in phosphate buffered saline (PBS) for >1 hour, rinsed 6x in PBS, and incubated for 5 nights in 1 μg/mL streptavidin conjugated to Alexa Fluor-568 (Invitrogen) in PBS that was supplemented with 1% Triton x-100 and 0.5% dimethylsulfoxide (DMSO) at 4°C. Following streptavidin incubation, tissue was washed 6x in PBS and mounted and coverslipped with Vectashield HardSet (Vector Labs). Filled RGCs were typically imaged immediately post-recording. Filled neurons were imaged on a 2-photon microscope (Scientifica, Uckfield, UK) with the Mai Tai Ti:sapphire laser (Spectra-Physics, Santa Clara, CA) tuned to 800 nm. Four images per plane were acquired along the z-axis at 1 micron spacing. For analysis, each image in the plane was averaged in ImageJ and dendrites, axons, and somata were traced using the Simple Neurite Tracer plug-in (Longair et al., 2011).

The morphological class of each filled TC neuron was identified using a previously published approach (Krahe et al., 2011) that quantitatively measures dendritic field asymmetry by performing a Sholl analysis with five concentric Sholl rings (Sholl diameter = 1/5 of dendritic field diameter) on four quadrants of the field. This allowed for calculation of an index of dendritic orientation (DOi), which equals the ratio of the intersections summed from opposite quadrants. We used thresholds for DOi = 0.0-0.5 for X-cells, 0.51-0.7 for W-cells, and 0.71-1.0 for Y-cells. The majority of our filled cells were from the ventromedial dLGN and had Y-cell morphology by these criteria, consistent with published assessments of dLGN regional preference (Krahe et al., 2011). No X-cells were detected in our sample and the few W-cells we did fill were excluded from further analysis. Dendritic Sholl analysis was performed using the Sholl analysis ImageJ plug-in (excluding TC neuron axon and somata) (Ferreira et al., 2014) while dendritic field area was calculated as the area of a convex polygon drawn by connecting dendritic tips in a 2D projection of the filled cell.

### Immunofluorescence staining

To measure RGC density, we performed immunofluorescence labeling of RGCs using two approaches. To label ChR2-EYFP expressing RGCs, an anti-GFP antibody, which is also sensitive to EYFP, was used to label ChR2-expressing RGCs while position in the retina was determined using an antibody sensitive to S-cone opsin (Sondereker et al., 2018), which is expressed in a gradient along the dorsal-ventral axis of the retina (Applebury et al., 2000). To label all RGCs, we used a guinea pig-anti-RBPMS antibody (Rodriguez et al., 2014). Retinas were dissected in aCSF bubbled with CO_2_/O_2_, mounted on nitrocellulose membrane (Type AAWB, 0.8 micron pore size, Millipore, Burlington, MA) and fixed by immersion in 4% paraformaldehyde in PBS for ~1 hour. Retinas were washed, blocked and permeabilized in a solution of 1% Triton X-100, 0.5% DMSO and 5.5% donkey serum for one hour before being incubated in primary antibodies for 2-4 days at 4°C. Since we used a goat-raised secondary for visualizing RBPMS-stained RGCs, we also included 5.5% goat serum when performing RBPMS labeling. Following primary antibody incubation, retinas were washed six times, blocked/permeabilized again, incubated in AlexaFluor-conjugated secondary antibodies (1:200) for 2 hours at room temperature, washed again, and mounted and coverslipped on SuperFrost plus slides with VectaShield Hardset. Labeled RGCs were imaged in central retina (~500 microns from the optic nerve head) and peripheral retina (~1700 microns from optic nerve head) in four quadrants (temporal, nasal, dorsal, ventral) identified based on S-opsin gradient. Imaging was performed with a Scientifica 2-photon microscope (as above), and RGCs were manually counted using the Cell Counter ImageJ plugin by graders blinded to treatment condition.

For immunofluorescence in dLGN slices, mice were euthanized and brains were dissected, as above. Brains were rinsed briefly in PBS, and immersed in 4% paraformaldehyde for 2 nights at 4°C. They were then rinsed in PBS, cryoprotected in 30% sucrose, embedded in 3% agarose, and cut into 50 micron sections on a vibrating microtome (Leica Biosystems VT1000S, Buffalo Grove, IL). Slices were collected immediately post-cutting on SuperFrost Plus slides (ThermoFisher) and allowed to dry before storage at −20°C. Immunohistochemistry was performed on mounted dLGN slices following the same protocol for retina except that the DMSO was omitted from the blocking/permeabilization solution and the Triton X-100 concentration was reduced to 0.5%. Control and OHT tissue was processed in parallel. Antibodies used in this study are listed in Table 1. dLGN imaging was performed on a 2-photon microscope with a 20x water-immersion objective (Scientifica) with the laser tuned to 800 nm. The size of the imaged region, laser power, and detector sensitivity was identical for all acquired images. For quantification of RGC axon terminal density and size, the vGlut2 signal was automatically thresholded and puncta were detected and measured using the Synapse Counter plug-in in ImageJ (Dzyubenko et al., 2016).

**Table 1:**
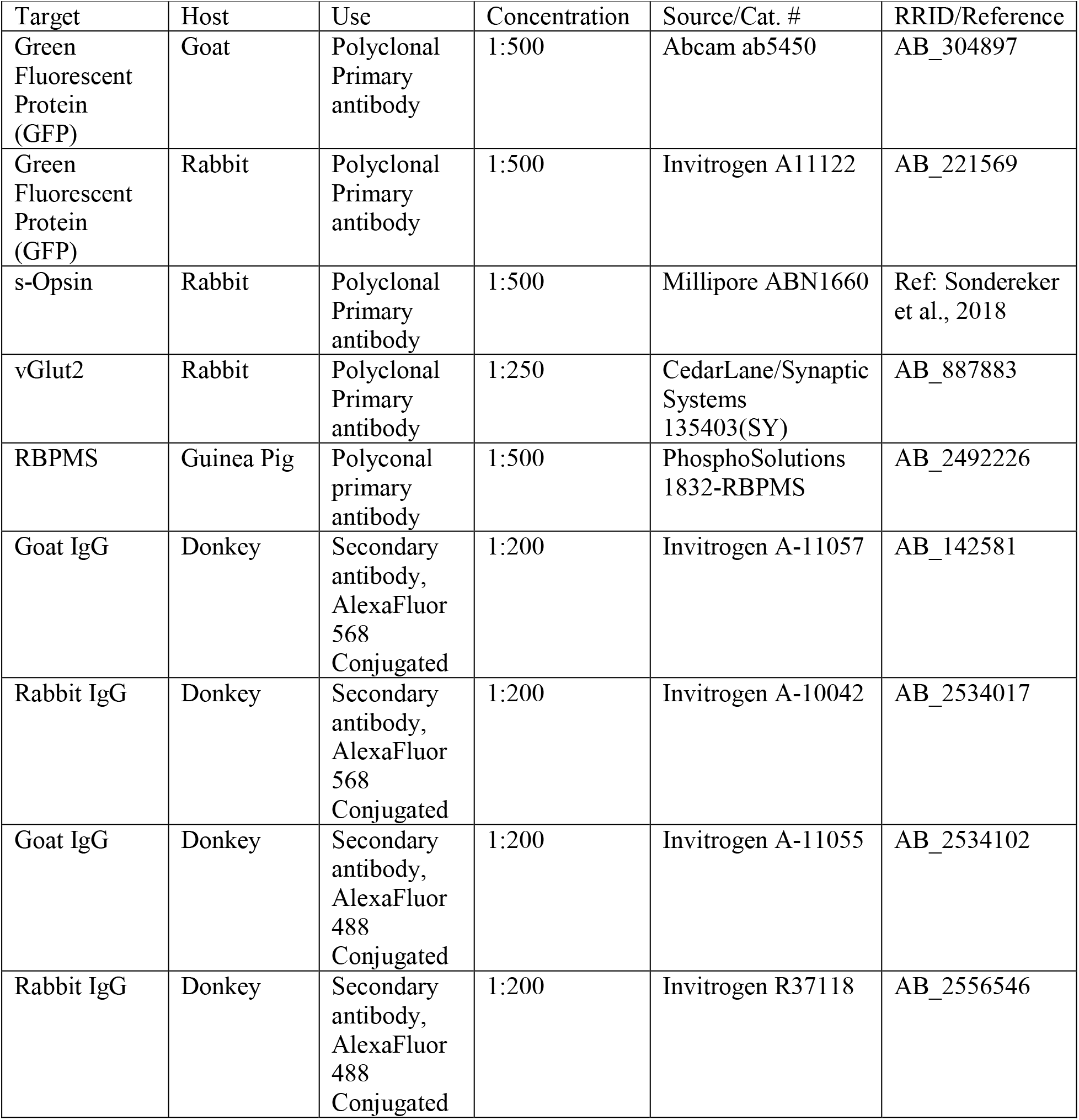
Antibodies used in this study. Host species, target, concentration, source and RRID/reference information for primary and secondary antibodies used for immunofluorescence experiments.

### Analysis and statistics

Unless noted otherwise, data are presented as mean ± SEM. When applicable, sample sizes indicate the number of cells recorded and the number of mice. Statistical significance was typically assessed using an unpaired Student’s t-test, although we used paired t-tests for paired data sets and Komolgorov-Smirnov tests to assess differences in population distributions of data, as indicated. Clampfit 10, Microsoft Excel, and GraphPad Prism 7 were used for statistical analyses.

## Results

To induce a modest ocular hypertension in mice, we performed bilateral injections of 10 micron polystyrene microsphere into the anterior chamber. Microbead injection caused a sustained increase in IOP of 4.8 ± 0.2 mmHg (n = 67 eyes) over baseline at 4 week post-injection (36 ± 3% increase). These values were significantly different from saline-injected controls (0.39 ± 0.15 mmHg over baseline, p=5E-30; 3 ± 1% increase over baseline, p=1.4*10^−19^; n = 69 eyes; Figure 1A). This modest OHT mimics OHT in human patients with primary open angle glaucoma and contrasts with the large and/or transient OHT used in some animal studies (Liu et al., 2017; Ly et al., 2011; Ou et al., 2016; Yücel et al., 2001).

**Figure 1.**
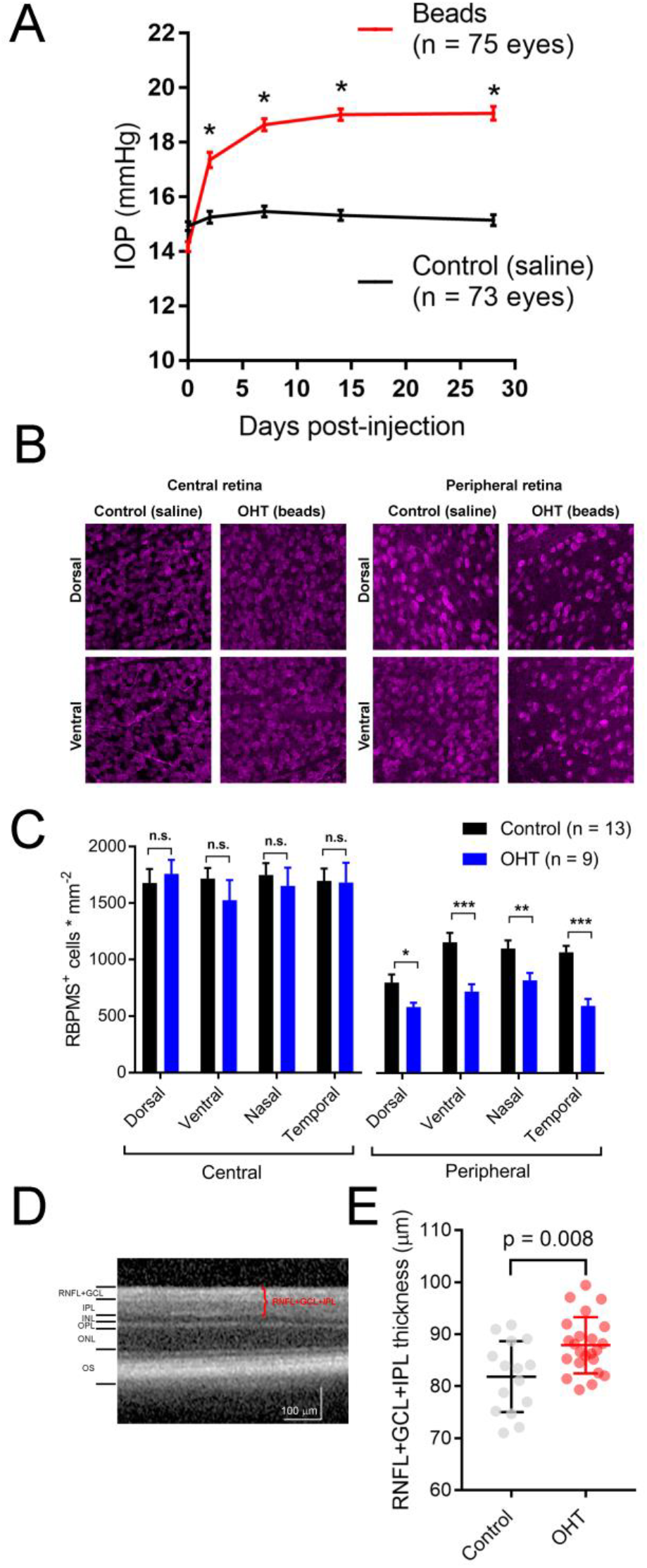
Ocular hypertension leads to retinal ganglion cell loss and inner retinal pathology. A) Anterior chamber injection of 10-micron microspheres led to an increase in intraocular pressure (IOP) while injection of saline did not. *p<<0.005. B) 2-photon image of retinal ganglion cells labeled with an anti-RBPMS antibody. Images from ventral and dorsal quadrants are in central and peripheral retina (~500 and ~1700 μm from optic nerve head) from saline- and bead-injected mice are shown. C) Quantification of RBPMS-stained RGC density in central and peripheral retina showing lower RGC density in peripheral, but not central, retina in bead-injected mice. *p<0.05; **p<0.01; ***p<0.001. D) Optical coherence tomography image showing identification of retinal layers. The RNFL+GCL+IPL thickness was measured. E) Quantification of OCT imaging (measured at 500 μm from Bergmeister’s Papilla), shows a modest inner retinal thickening in ocular hypertensive mice. The RNFL+GCL+IPL thickness is plotted individually for each eye and the error bars are standard deviation.

Five weeks post-injection, we harvested eyes and measured RGC density in four quadrants of central and peripheral retina by immunofluorescence staining of RGC somata with an anti-RBPMS antibody and two-photon imaging. Following counting of RGC somata, we found that RBPMS-stained RGC density was similar in all four quadrants of central retinas from control and OHT mice, but was significantly reduced in all four quadrants of peripheral retina (Figure 1B & C). This is consistent with other studies in which peripherally-located RGCs appear to degenerate at an earlier time point than those located in the central retina (Chen et al., 2011).

Optical coherence tomography (OCT) is an imaging technique commonly used in clinical settings to assess ocular disease progression *in vivo*. We performed OCT imaging of OHT mice and saline-injected control mice to test for signs of inner retinal pathology in central retinal layers (RNFL+GCL+IPL). 5 weeks post-injection, we found a small, but statistically significant thickening of the inner retinal layers (Figure 1D & E). In saline-injected control mice, RNFL+GCL+IPL thickness was 81.8 ± 1.8 μm (n = 15 eyes), while it was 86.9 ± 1.1 μm in bead-injected ocular hypertensive eyes (n = 23, p = 0.008). This thickening, which might be the result of axonal swelling due to inhibited axoplasmic transport (Abbott et al., 2014), confirms that microbead OHT leads to detectable changes in retinal structure.

Having found that microbead injection leads to a modest and sustained OHT in addition to RGC degeneration and other signs on retinal pathology, we next sought to determine how OHT affects RGCs and their output synapses in the dLGN. Some current evidence suggests that different RGC classes are differentially affected by OHT (Della Santina et al., 2013; El-Danaf and Huberman, 2015; Ou et al., 2016; Risner et al., 2018). Therefore, we opted to focus on a somewhat narrow subpopulation of RGCs selectively expressing fluorescent and optogenetic reporters in retinal ganglion cells that express the photopigment melanopsin. Melanopsin-expressing RGCs are best known for their roles in irradiance encoding and projections to non-image-forming brain nuclei such as the suprachiasmatic nucleus or olivary pretectum where they regulate reflexive responses to light (i.e., pupillary constriction and circadian photoentrainment) (Berson et al., 2002; Hattar et al., 2002, 2006). However, some melanopsin-expressing RGCs express only low levels of melanopsin protein, receive strong rod- and cone-driven synaptic inputs, send axonal projections to the dLGN, and underlie conscious, image-forming vision. M4-type melanopsin RGCs are identical to the population of On-sustained αRGCs and are a prime example of this phenomenon (Estevez et al., 2012).

We first verified that OHT produces detectable effects on the structure and function of M4 cells by targeting them for cell-attached patch clamp recording and whole-cell dye filling in retinas from *Opn4*-cre/+;Z/EG mice. We found that OHT was associated with a lower spontaneous spiking frequency relative to controls (control = 24 ± 2 Hz, n = 15; OHT = 11 ± 1 Hz, n = 19; p = 0.0002). Sholl analysis of dye-filled and reconstructed dendrites revealed a similar Sholl profile to a previous study of M4 RGCs (Estevez et al., 2012). In retinas from ocular hypertensive mice, the peak number of Sholl crossings was reduced, indicative of a slight reduction in dendritic complexity (Figure 2D-F).

**Figure 2.**
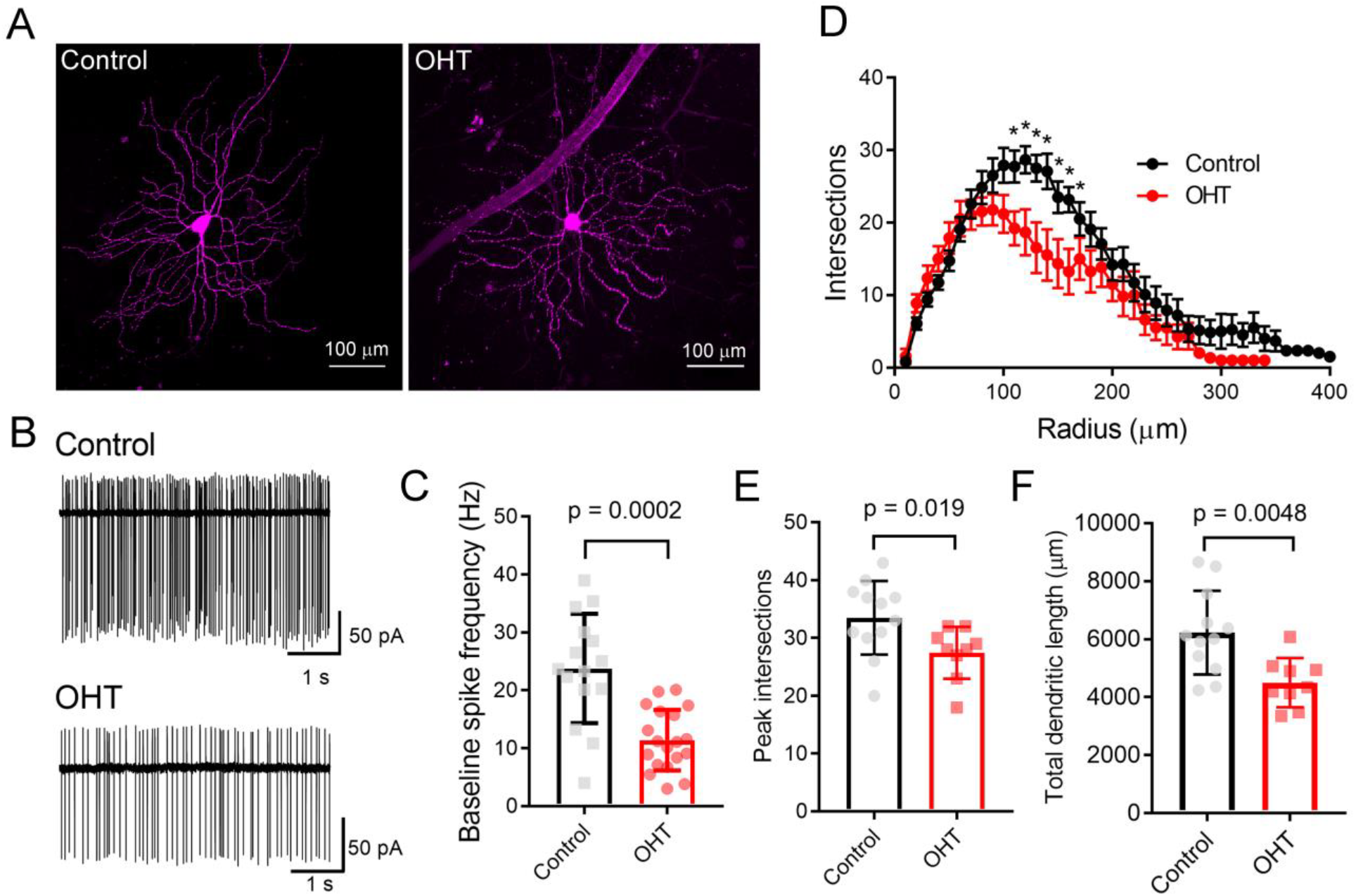
Ocular hypertension alters the structure and function of M4-type RGCs. A) Maximum intensity projection images of dye-filled M4/On-αRGCs filled with CF568 dye during whole cell recording in flat mount retinas. B&C) In cell-attached measurements of spontaneous spiking (in the dark, in the absence of stimulus), the spontaneous spike frequency was lower in OHT than control. D) Sholl analysis of M4 dendrites shows that dendritic complexity was slightly reduced relative to control cells. E) Comparison of peak number of Sholl intersections in control and OHT M4 RGCs. F) Total dendritic length was reduced in M4 RGCs from OHT retinas. Values for each cell are plotted individually in C, E, and F and the bar graphs represent mean ± standard deviation.

We next sought to determine how OHT affects the RGC output to the dLGN by using ChR2 expression to selectively activate melanopsin-expressing RGC axons in acute dLGN brain slices from *Opn4*-cre/+;Ai32 mice. We prepared 250 micron coronal brain slices through the dLGN and targeted TC neurons for whole-cell voltage clamp recording (Figure 3). Using the protected recovery method of brain slice preparation (Ting et al., 2014, 2018), we could routinely find an abundance of healthy cell bodies in slices from adult animals (>postnatal day 100). Stimulation with a pair of 460 nm LED flashes (0.5 ms, 200 ms interval) evoked robust inward currents that displayed paired-pulse synaptic depression (Figure 3B), which is characteristic of retinogenciulate synaptic transmission (Chen and Regehr, 2000). γDGG (200 μM) and cyclothiazide (100 μM), which prevent AMPA receptor saturation and desensitization, respectively, together enhanced current amplitude and slowed decay kinetics; amplitude was increased by 42 ± 6% (n = 13, p = 0.0002, paired t-test) while the decay time constant was slowed from 2.9 ± 0.2 ms to 7.9 ± 0.6 ms (n = 13, p = 0.00000009, paired t-test). Additionally, the paired pulse ratio was increased from 0.52 ± 0.1 to 0.62 ± 0.1 (n = 13, p = 0.0015), indicating that AMPA receptor desensitization/saturation contribute to short-term synaptic depression at this synapse (Budisantoso et al., 2012). Additional pharmacology and current-voltage experiments (Figure 3C-G) confirmed that these responses were mediated by excitatory glutamatergic synaptic transmission; responses reversed near the cationic equilibrium potential (7.4 ± 1.1 mV, n = 21) and were blocked by 40 μM CNQX (98 ± 0.3 % reduction in amplitude, n = 7, p= 6*10^−13^, one-sample t-test). Slower NMDA-receptor currents could be detected when TC neurons were depolarized to +40 mV (Figure 3F&G). The AMPA/NMDA ratio was similar to that recorded with electrical stimulation of the optic tract in separate parasagittal dLGN slice experiments (ChR2 stimulation AMPA/NMDA = 3.2 ± 0.3, n = 9; electrical stimulation AMPA/NMDA = 3.1 ± 0.5, n = 5; p = 0.844). EPSC responses to ChR2 stimulation were also blocked by 1-2 μM tetrodotoxin (Figure 3H; 98 ± 2% reduction in amplitude, n = 7, p = 1.4*10^−11^, one-sample t-test) or 200 μM Cd^2+^ (Figure 3I; 99 ± 0.3% reduction in amplitude, n = 4, p = 0.00000004, one-sample t-test), indicating they depended on presynaptic action potential spiking and Ca^2+^ entry.

**Figure 3.**
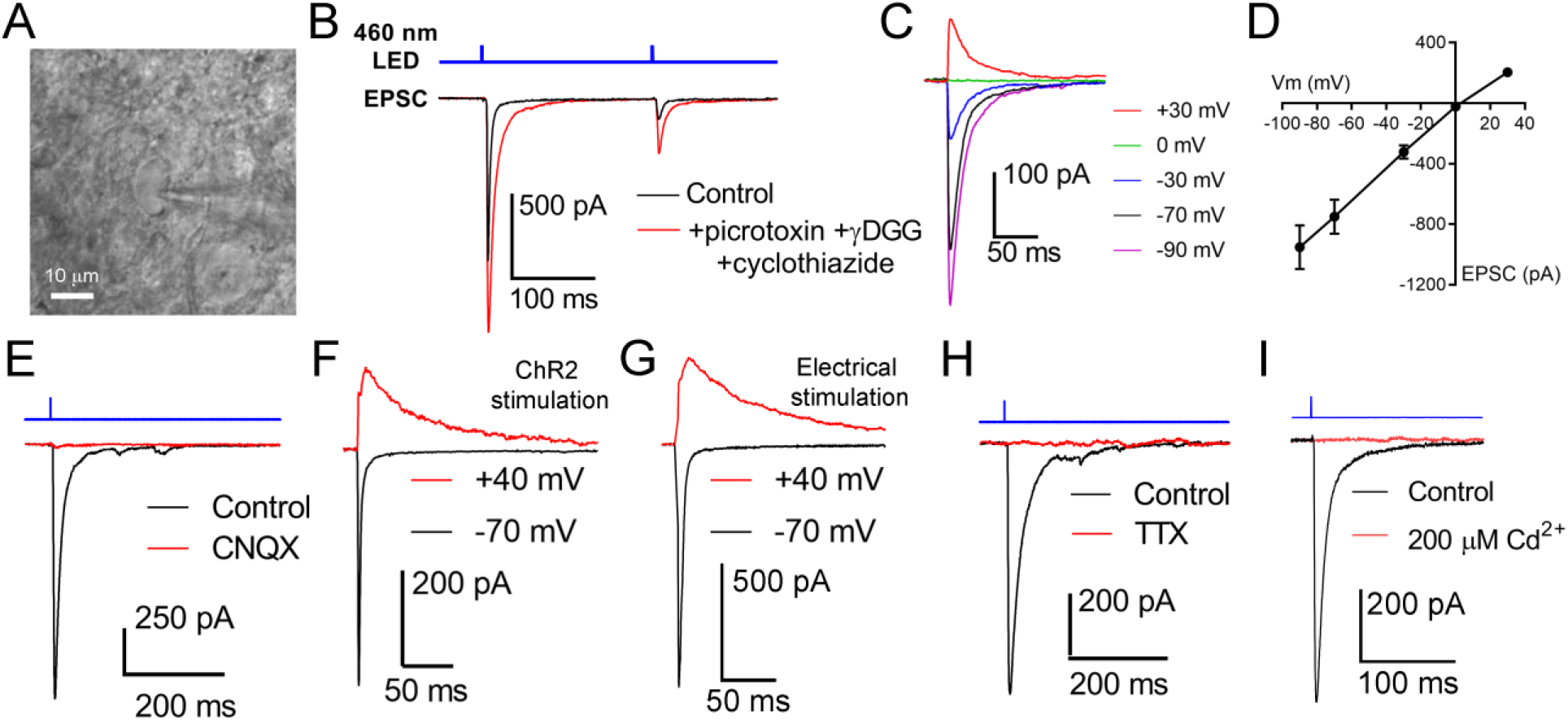
Activation of channelrhodopsin in dLGN brain slices evokes post-synaptic responses in TC neurons. Thalamocortical neurons were targeted for whole-cell patch-clamp recording in dLGN brain slices (A). B) Stimulation with a pair of full strength flashes from the TLED (0.5 ms duration, 200 ms interval, 1.5 mW) evoked large currents displaying paired pulse depression. Current amplitude was enhanced and decay kinetics slowed by addition of 200 μM γDGG and 100 μM cyclothiazide. Picrotoxin (60 μM) was added to block inhibition. C & D) In a current-voltage experiment, currents reversed near the cationic equilibrium potential (~0 mV). E) ChR2-evoked currents were sensitive to the AMPA receptor blocker CNQX, consistent with responses being excitatory post-synaptic currents (EPSC). F) Slower, presyumptive NMDA receptor-mediated EPSCs could be detected at depolarized potentials. G) The AMPA/NMDA ratio was similar when the optic tract was stimulated using an extracellular electrode in parasagittal brain slices. H) EPSCs were blocked by the Na^+^ channel blocked Tetrodotoxin (1-2 μM). I) EPSCs were blocked by the Ca^2+^-channel blocked Cd^2+^ (200 μM).

Other neurosensory disorders, such as conductive hearing loss, are associated with changes in synaptic vesicle release probability (Zhuang et al., 2017). To test for changes in synaptic vesicle release probability (Pr) associated with OHT, we varied the interval between a pair of pulses from 100 ms to 2 s while measuring the paired pulse ratio (PPR) of the evoked EPSCs (PPR = EPSC2/EPSC1) in the presence of γDGG and cyclothiazide to prevent AMPA receptor saturation and desensitization, respectively (Sakaba et al., 2002) (Figure 4). As is typical for synapses with a high Pr, the PPR increased with increasing stimulus interval, which is likely attributable to synaptic vesicle pool refilling. The PPR measured with ChR2 stimulation was similar to the PPR measured using electrical stimulation of the optic tract in separate parasagittal dLGN slice recordings at these intervals (n = 7 cells from 4 mice; Figure 4C&D). In brain slices from OHT mice, the PPR measured with ChR2 stimulation was significantly reduced at most intervals tested; at a 100 ms interval, for instance, the PPR was lower in OHT mice (0.44 ± 0.04; n = 15 cells from 11 mice) compared to controls (0.66 ± 0.04, n = 21 cells from 13 mice; p = 0.0005). Given a long enough time interval (i.e. 60 seconds), the PPR recovered to ~1 in both control and OHT conditions. This decrease in the PPR (stronger synaptic depression) in bead-injected mice likely indicates that OHT triggers an increase in presynaptic vesicle release probability (Pr). Consistent with prior studies of retinogeniculate synapses (Chen and Regehr, 2000; Lin et al., 2014; Litvina and Chen, 2017), there was substantial variability in the EPSC amplitudes recorded in TC neurons. This phenomenon is a combination of both the variability in the strength of the inputs from single RGC axons (the single-fiber amplitude), which can range from 10 pA to >3 nA, and variability in the degree of RGC convergence onto single TC neurons (Litvina and Chen, 2017). However, despite the increase in Pr we did not observe any increase in the EPSC amplitude measured in response to a maximal LED stimulus (1.5 mW; Figure 4G&H).

**Figure 4.**
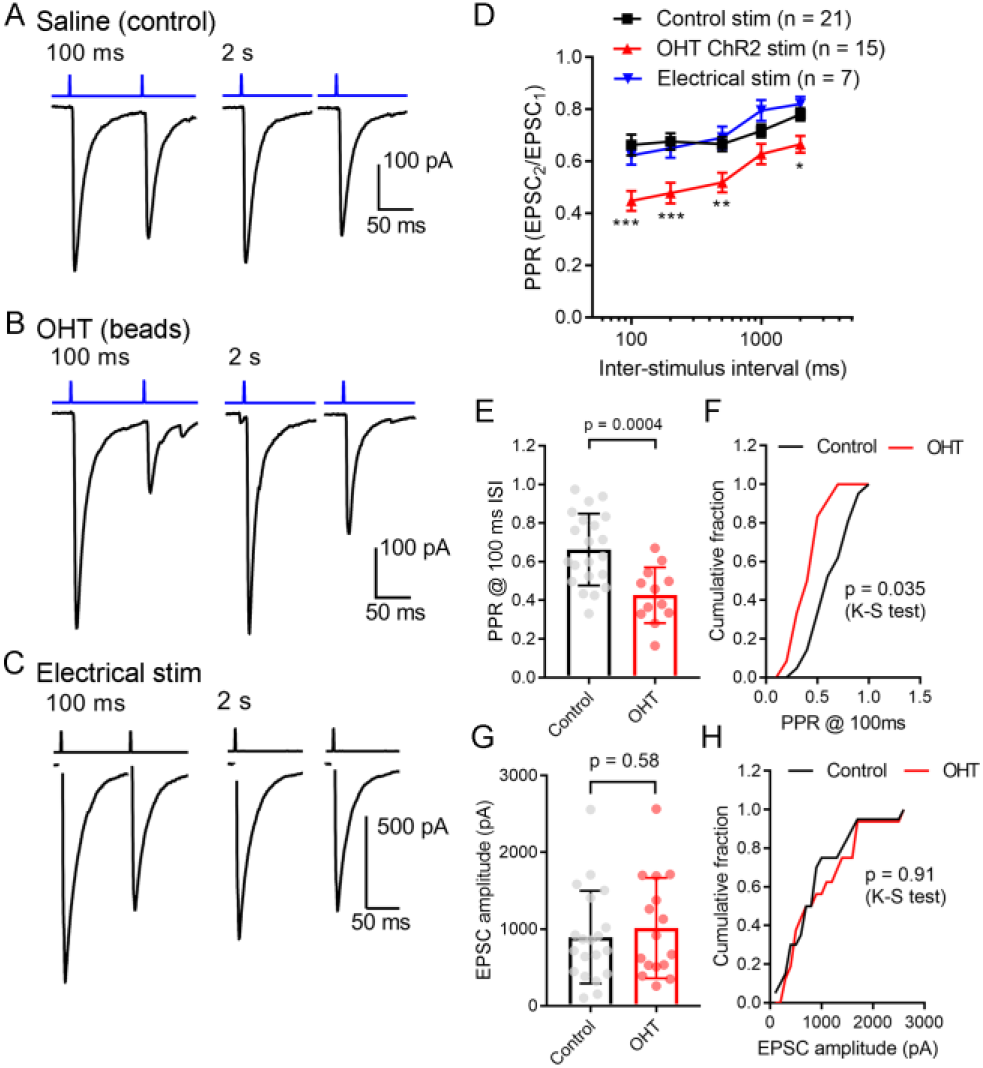
Ocular hypertension increases synaptic depression at retinogeniculate synapses. A) In brain slices from control animals, stimulation with a pair of 0.5 ms 460 nm LED flashes delivered through the microscope objective triggered post-synaptic currents displaying synaptic depression. In OHT (B), depression was more pronounced. C) Paired pulse experiment in which the optic tract was stimulated with an extracellular electrode in a parasagittal brain slice through the dLGN. The PPR was similar to the ChR2-evoked response in brain slices from saline-injected (control mice). The stimulus artifact was deleted for display. D) The paired pulse ratio (EPSC2/EPSC1) plotted against stimulus interval showed that although PPR increased with increasing stimulus interval, it was lower in bead-injected OHT mice than in controls. PPR values recorded in response to electrical stimulation of the optic tract in parasagittal slices are shown for comparison. Unpaired t-test was used to compare PPR in from control and OHT mouse brain slices. **p<0.005, ***p<0.001, *p<0.01. E & F) The paired pulse ratio measured at a 100-ms interval was lower in OHT brain slices relative to controls. G&H) There was no significant difference in the EPSC amplitude evoked by a saturating (1.5 mW) LED stimulus. Values are plotted for each individual cell in E&G and the bar graphs represent mean ± standard deviation.

As a means of performing a more quantitative measurement of vesicle release probability in control vs. OHT retinogeniculate synapses, we next activated RGC axons with a relatively high-frequency train stimulus (10 Hz, 30 stimuli) (Figure 5) (Sakaba et al., 2002). In this analysis, we plotted the cumulative EPSC amplitudes and fit the linear portion (pulses 15-30) with a straight line extrapolated to the vertical axis. The ratio of the first EPSC to the y-intercept is Pr. In control brain slices, the Pr measured using 10 Hz activation of ChR2 was 0.39 ± 0.04 (n = 21 cells from 12 mice), which was not significantly different from Pr measured in an identical fashion from EPSCs evoked with 10 Hz electrical stimulation of the optic tract in parasagittal slices (Figure 5C&D; Pr = 0.35 ± 0.02; n = 8 cells from 4 mice; p = 0.3). In experiments from OHT mice, Pr was significantly enhanced to 0.54 ± 0.03 in OHT (n = 13 cells from 10 mice, p = 0.005; Figure 5D&E), consistent with the reduced PPR described above (Figure 4).

**Figure 5.**
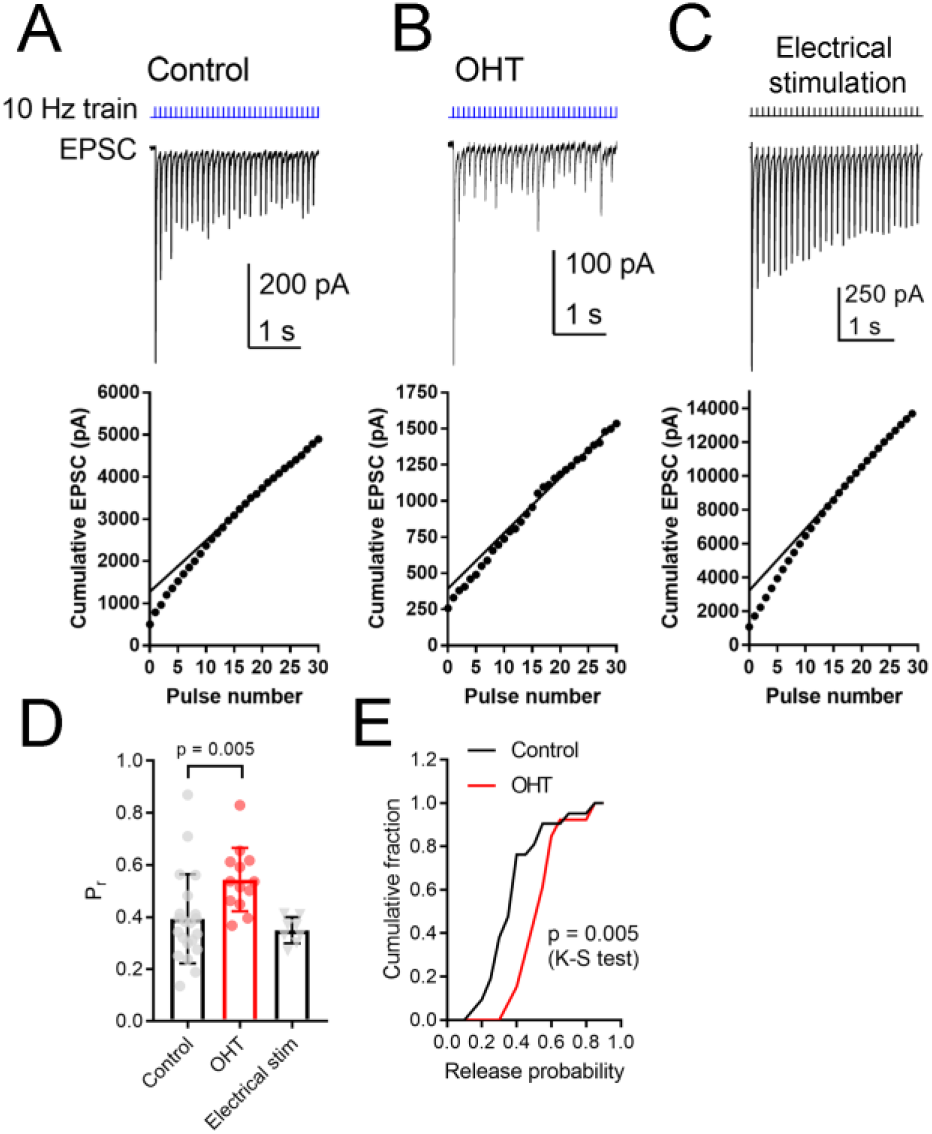
OHT increases presynaptic vesicle release probability at retinogeniculate synapses. A train of LED stimuli (10 Hz, 30 stimuli) was used to assess synaptic vesicle release probability at RGC synapses onto TC neurons in control (A) and OHT (B) dLGN slices. Upper panels show the EPSC and lower panels show the cumulative EPSC with a straight line fit to stimuli 15-30 and extrapolated to the vertical axis. The ratio of the amplitude of the first EPSC to the vertical intercept of the fit is the release probability. C) Responses to 10 Hz electrical stimuli delivered to the optic tract in parasagittal brain slices were similar to responses evoked by ChR2 stimulation in brain slices from saline-injected (control) mice. D) The release probability (Pr) was elevated in recordings from OHT mice relative to recordings from saline-injected controls. The Pr measured from extracellular optic tract stimulation in parasagittal slices (“Electrical stim”) was similar to control values obtained with ChR2 stimulation. Values for each recorded cell are plotted individually and the bar graphs display mean ± standard deviation. E) Cumulative probability distribution of Pr values in control and OHT.

To test whether OHT leads to changes in ChR2 expression or excitability of RGC axons, we next recorded ChR2-driven population spikes from RGC axons in the optic tract using a range of 460 nm LED stimulus intensities (Figure 6). The half-maximal stimulus intensity (I_50_) values for the population spike recorded using an extracellular electrode positioned in the optic tract dorsal to the dLGN were similar (control = 68 ± 28 μW, n = 3; OHT = 58 ± 13 μW, n = 3; p = 0.62). These data suggest that early-stage OHT does not affect the ability of RGC axons to spike following ChR2 activation in our brain slice preparation.

**Figure 6.**
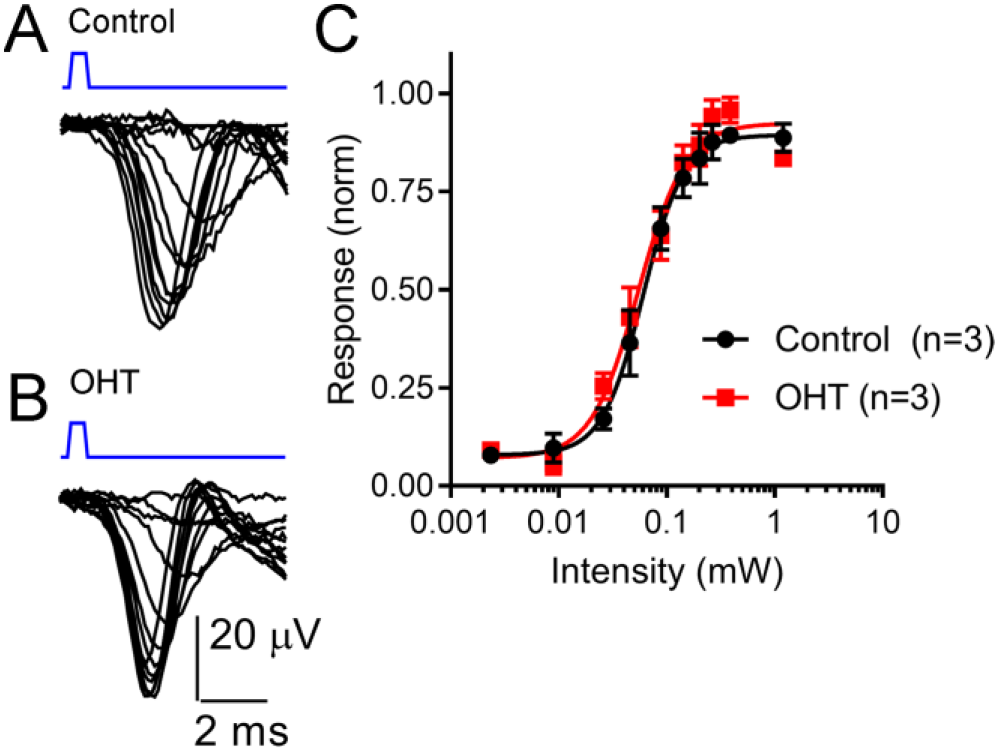
Sensitivity of the optic tract to LED stimulation of ChR2 was not altered by OHT. A&B) An extracellular electrode was positioned in the optic tract in coronal sections from saline-injected (control) or OHT mice in order to record population spiking in response to ChR2 activation with a series of LED stimulus intensities (2 μW - 1.2 mW). Each trace is an average of 3-10 individual responses at each intensity. Stimulus timing (0.5 ms LED flash) is marked above the traces in blue. C) A plot of normalized response amplitudes shows that optic tract sensitivity to ChR2 was not altered by OHT.

We next used two approaches to monitor the contribution of single presynaptic vesicles (quantal amplitude) to the multiquantal EPSC (Figure 7). We first recorded spontaneous miniature EPSCs (mEPSCs) in the absence of stimulation (Figure 7A-E). In these recordings, the mEPSC amplitude was not significantly different between control and OHT (control: 9.2 ± 0.9 pA, n = 8 cells from 5 mice; OHT: 8.8 ± 0.8 pA, n = 7 cells from 6 mice; p = 0.76, t-test). Likewise, when we compared the cumulative distributions of 50 events from each recorded cell, the amplitude distributions were not significantly different between control and OHT (p = 0.46, K-S test). We also compared the frequencies of mEPSCs in the absence of stimulation (Figure 7D&E), finding that the mean mEPSC frequency was lower in TC neurons from OHT mice (9.3 ± 1.7 Hz, n = 7 cells from 6 mice) compared to TC neurons from control mice (17.9 ± 3.2 Hz, n = 8 cells from 5 mice, p = 0.038). Likewise, when we compared the distributions of inter-event intervals for 50 events per recorded cell, we found that the cumulative distribution was significantly shifted to longer intervals (p<0.00001, K-S test). This indicates that OHT reduces the spontaneous synaptic input to dLGN TC neurons.

**Figure 7.**
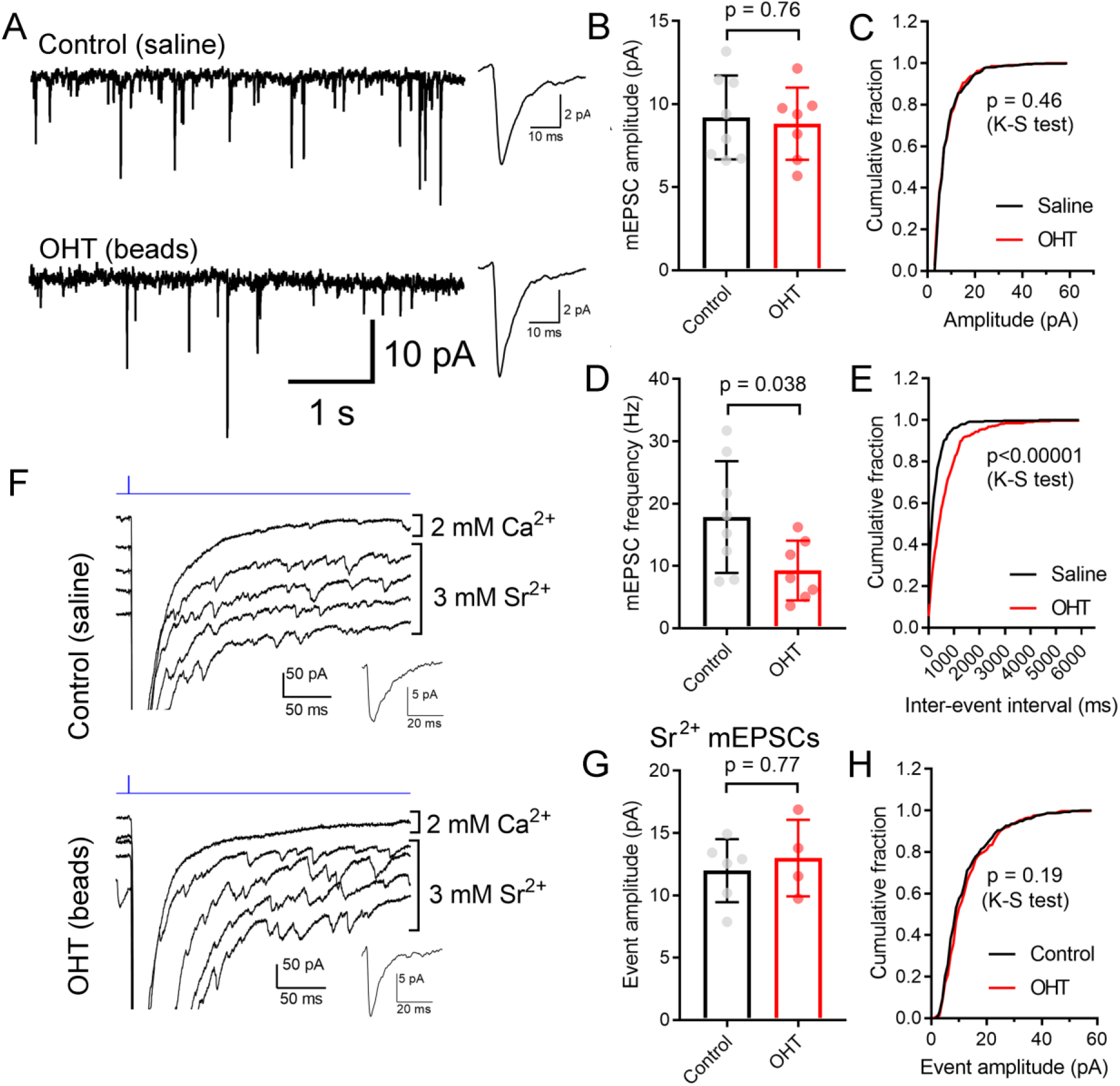
Influence of OHT on single-vesicle synaptic currents in TC neurons in the dLGN. A) mEPSCs recorded in the dLGN of TC cells in the absence of stimulation. These TC cells displayed evoked multiquantal EPSCs in response to ChR2 stimulation with 460 nm stimulation. Insets show mean mEPSC waveforms 100 individual mEPSCs detected in these recordings. B&C) The mEPSC amplitude was not significantly different between control and OHT TC neurons. D&E) The mEPSC frequency was significantly lower in OHT recordings whether assessed with an unpaired t-test (D) or with a Komolgorov-Smirnov test used to compare the cumulative distributions of 100 mEPSC inter-event intervals per recorded cell (E). F) To record properties of single-vesicle events evoked by ChR2 stimulation, CaCl_2_ in the aCSF was replaced with 3 mM SrCl_2_. A single trace recorded in with Ca^2+^ and four traces recorded with Sr^2+^ are shown. Insets show waveforms of detected mEPSC events. G&H) The amplitude of single-vesicle events evoked by ChR2 stimulation in Sr^2+^-containing aCSF were not significantly different between recordings from control and OHT mice. Values are plotted from individual cells in panels B, D, and G and the bar graphs show mean ± standard deviation.

mEPSCs recorded in the absence of stimulation represent a mix of glutamate release from ChR2-expressing RGC axon terminals, RGC axon terminals that do not express ChR2, and excitatory inputs arising from the corticothalamic tract. To examine just the single-vesicle events arising from ChR2 stimulation of the optic tract, we replaced the extracellular CaCl_2_ in the aCSF with 3 mM SrCl_2_ (Figure 7F-H). This de-synchronizes synaptic vesicle release, allowing us to resolve single-vesicle release events following optic tract stimulation. In these experiments (conducted in the presence of 200 μM γDGG and 100 μM cyclothiazide), single-vesicle EPSCs were not different between control (12.0 ± 1.0 pA, n = 6 cells from 4 mice) and OHT (13 ± 1.5 pA, n = 4 cells from 3 mice) whether compared using a t-test (p = 0.61) or a K-S test to compare cumulative distributions of 75 events per cell (p = 0.19). Thus, mEPSC amplitudes appear unchanged by OHT.

Prior studies have suggested that RGC axon terminals in the dLGN and superior colliculus persist fairly late into disease in rodent glaucoma models (Crish et al., 2010; Smith et al., 2016). However, to test whether this reduction in mEPSC frequency might be due to the loss or atrophy of RGC axon terminals, we stained 50-micron dLGN sections from control or OHT mice using an antibody sensitive to vGlut2, which is a selective marker of RGC axon terminals in the dLGN (Fujiyama et al., 2003; Land et al., 2004), and imaged vGlut2 puncta in the ventromedial dLGN on a 2-photon microscope (Figure 8). Automated detection and analysis of vGlut2 puncta revealed that there was no significant difference in vGlut2 density between control and OHT (control: 3.6 ± 0.4 puncta/1000 μm^2^, n = 11 brain slices from 11 mice; OHT: 3.4 ± 0.7 puncta/1000 μm^2^, n = 10 brain slices from 10 mice; p = 0.39; Figure 8B). Additionally, vGLut2 punctum size did not differ between control and OHT whether we compared average punctum size per brain slice (control = 26.8 ± 2.0 μm^2^; OHT: 27.1 ± 0.18 μm^2^; p = 0.69; Figure 8C) or the cumulative distribution of the size of 300 puncta per brain slice (p = 0.86, K-S test; Figure 8D).

**Figure 8.**
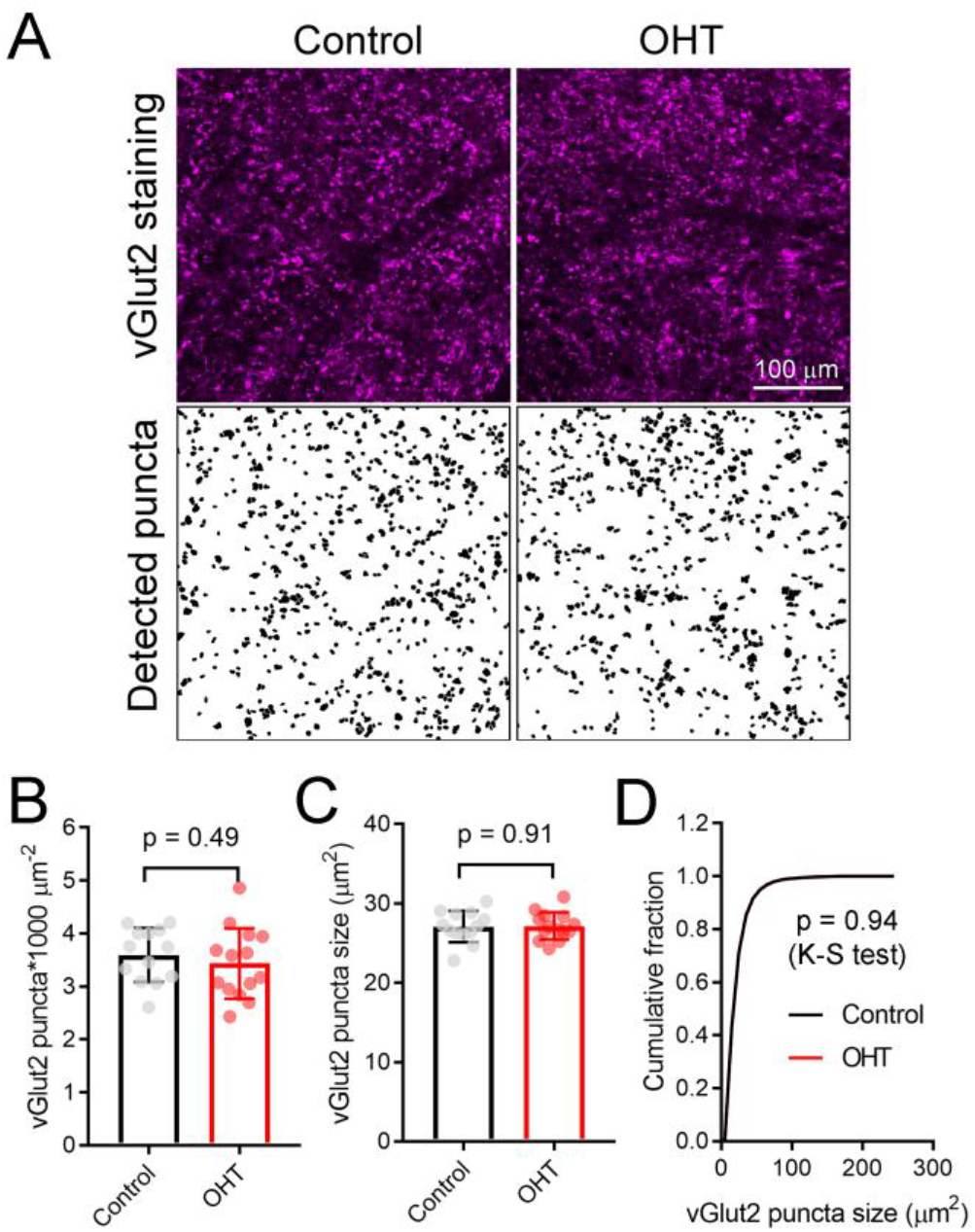
Five weeks modest OHT does not alter vGlut2 staining in the dLGN. A) 2-photon images of a 1.5 micron optical section of dLGN stained for vGLut2, which marks RGC axon terminals in control and OHT conditions. Lower panels show vGlut2 puncta that were automatically detected in ImageJ. B-D) Neither the density nor the size of vGlut2 puncta differed between control and OHT dLGN. The density and size of puncta for each brain slice are plotted individually and the mean ± SD is plotted in the bar graph (B&C). D) The cumulative distribution of 300 detected vGlut2 puncta per brain slice was nearly identical for control and OHT.

In the course of whole-cell recording, we also filled TC neurons with neurobiotin in order to test for early signs of OHT-triggered loss of dendrites that might contribute to the detected reduction in mEPSC frequency (Figure 9). While later-stage dendritic changes have been reported in primate studies (Gupta et al., 2007; Liu et al., 2011; Ly et al., 2011), it is unclear whether this process occurs earlier in the course of the disease. After reacting with streptavidin and imaging on a 2-photon microscope, we reconstructed dendritic arbors of neurobiotin-filled TC neurons and analyzed cells with Y-cell morphology (Krahe et al., 2011) for Sholl analysis. Filled TC neurons in our sample were located in the medial dLGN core, a region that receives the bulk of innervation from RGCs labeled in the Opn4-Cre mouse (Ecker et al., 2010; Estevez et al., 2012; Stabio et al., 2018). When we analyzed dendritic complexity (Figure 9B&C), we found a small reduction in the peak number of Sholl intersections (at ~50-70 microns from the somata), from 37.5 ± 4.5 in control (n = 12) to 28.7 ± 2.2 in OHT (n = 9; p = 0.002). However, the total dendritic field area (Figure 9D) was not significantly different between control (8.2 + 0.6 10^4^ μm^2^) and OHT (8.0 + 0.6 10^4^ μm^2^) TC neurons (p = 0.85). This indicates that early-stage OHT leads to a modest loss of TC neuron dendrites located somewhat proximally to the cell body. As this region of TC neurons receives the bulk of RGC-originating driver inputs (in contrast to the more distal dendrites, which are sites for synaptic input from cortical origins) (Morgan et al., 2016; Rafols and Valverde, 1973), this likely underlies the observed reduction in mEPSC frequency in OHT and suggests that OHT leads to early-stage postsynaptic re-wiring of RGC connections with dLGN TC neurons.

**Figure 9.**
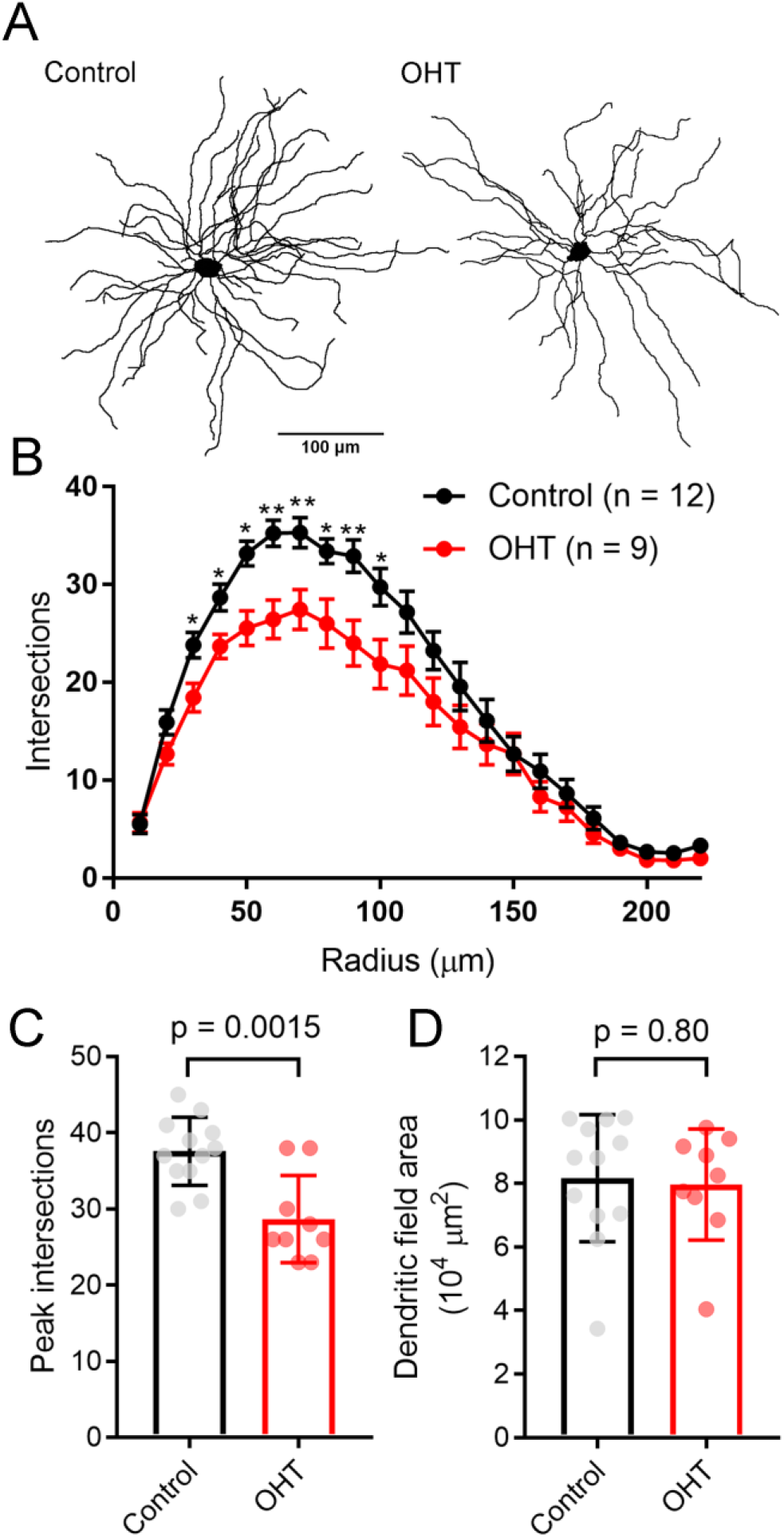
Dendritic complexity of TC neurons in the dLGN is reduced by OHT. A) dLGN TC neurons (Y-type profiles) were filled with neurobiotin during whole-cell recording for later 2-photon imaging and tracing of their dendritic profiles. B) Sholl analysis of TC neuron dendritic complexity shows a reduction in the number of dendrite crossings in OHT TC neurons, consistent with some dendritic pruning or retraction. *p<0.05 **p<0.01. C) The peak number of intersections was lower in TC neurons from OHT brain slices. D) The dendritic field area, measured by drawing a convex polygon contacting distal dendritic tips, was not significantly different between OHT and control groups. In C&D, vGlut2 density and puncta size are plotted individually from each brain slice and the bar graphs show mean ± standard deviation.

High IOP that eventually leads to glaucoma and blindness does so via degeneration of retinal ganglion cells (Calkins, 2012; Weinreb et al., 2014). The above findings demonstrate that a fairly modest and sustained OHT (~35% above baseline for 5 weeks) leads to detectable changes in the function of RGC synapses in the dLGN. Although we showed above that OHT leads to a detectable loss of RGCs in all four quadrants of the peripheral retina using a general RGC marker (RBPMS, Figure 2, above), it is not clear whether the population of RGCs providing dLGN input in our experiments were similarly affected, as melanopsin-expressing and On-type RGCs are generally believed to be more resistant to injury than other RGC populations (Cui et al., 2015; Della Santina and Ou, 2016; El-Danaf and Huberman, 2015; Ou et al., 2016; Valiente-Soriano et al., 2015b). To address this, we labeled ChR2-YFP expressing retinas from control and OHT *Opn4*-Cre/+;Ai32 mice using an antibody sensitive to GFP, which also labeled the YFP in the ChR2-YFP fusion protein (Figure 10). In contrast to the reduction of RGC density we observed with RBPMS-stained RGCs, there was no significant reduction in the density of ChR2-YFP RGCs either the center or periphery of all four quadrants of OHT retinas (Figure 10B). Thus, rather than following OHT-triggered degeneration of RGCs, these data indicate that the OHT-dependent changes in retinogeniculate synapses occur early in the disease process, prior to detectable loss of ChR2-expressing RGCs in the *Opn4*-cre/+;Ai32 mouse.

**Figure 10.**
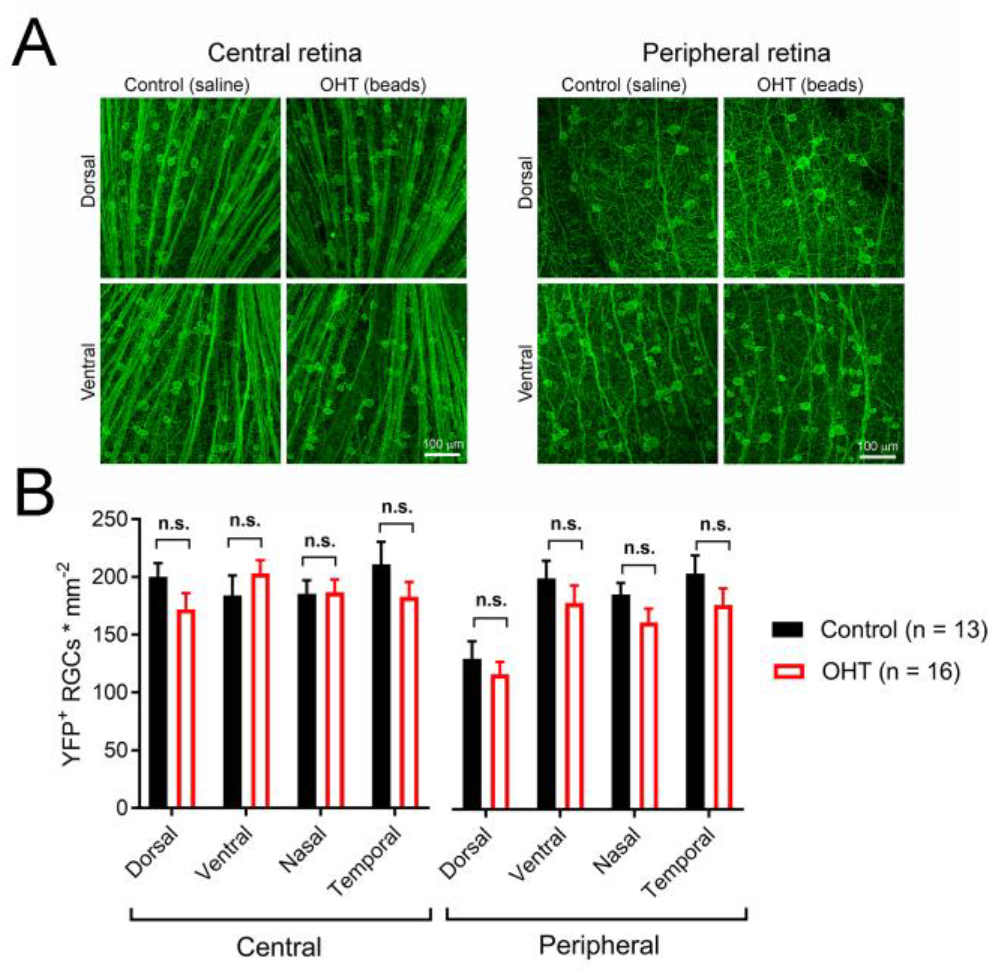
Five weeks of OHT does not lead to a detectable loss of melanopsin-expressing RGCs. A) In order to test for changes in the density of RGCs expressing the ChR2-YFP reporter, the somata, axons, and dendrites of ChR2-YFP expressing RGCs were stained using anti-GFP primary antibodies in retinal flat mounts. RGC somata were counted in central retina (~500 microns from optic nerve head) and peripheral retina (~1700 microns from optic nerve head) in dorsal, ventral, nasal, and temporal quadrants (determined by co-staining for S-opsin). B) YFP+ RGC somata density in four quadrants of central and peripheral retina. RGC density did not significantly differ between control and OHT groups (n.s., p>0.05).

## Discussion

The principal goal of this study was to test whether a moderate and sustained OHT that mimics human disease (Gazzard et al., 2003; Mao et al., 1991; Miglior and Bertuzzi, 2013) leads to changes in the visual pathway by quantitatively assessing the performance of RGC output synapses in the dLGN. Our core findings are that even the modest OHT triggered by bead injection alters presynaptic vesicle release probability without detectable effect on EPSC size. Additionally, the effects of OHT reach across the synapse to influence the dendritic structure of TC neurons. Notably, this occurred early in disease progression; synaptic changes were not associated with detectable loss of the ChR2-expressing RGC populations under study, although RGCs detected with a non-specific RGC marker were slightly reduced in the peripheral retina. These effects might therefore represent an early homeostatic responses to OHT-triggered effects on the optic nerve (Crish and Calkins, 2015; Risner et al., 2018). These novel findings demonstrate an early-stage impact of elevated eye pressure of the sort that mimics human disease on the function of the optic nerve and a key brain region receiving direct input from the retina and thereby shed light on a critical link between OHT and vision loss in glaucoma.

RGC axon damage at the optic nerve head is a key trigger for neurodegeneration in glaucoma and axonopathy is a core component of numerous neurodegenerative diseases (Salvadores et al., 2017). This has numerous impacts on the structure and function of RGCs including early dendritic pruning, changes in excitability, and loss of synaptic inputs (Della Santina et al., 2013; El-Danaf and Huberman, 2015; Ou et al., 2016; Pang et al., 2015; Risner et al., 2018). OHT also affects the optic nerve, triggering gliosis, axonal shrinkage, and diminished transport (Cooper et al., 2016; Crish et al., 2010, 2013; Martin et al., 2006; Pease et al., 2000; Quigley et al., 2000). Evidence from human patients and animal models has demonstrated clear changes in the structure and function of retinorecipient brain nuclei in glaucoma. One of the earliest effects of OHT measured in rodent glaucoma models is a deficit in axonal transport with the timeline of changes occurring in a distal-to-proximal order (Crish et al., 2010). Only much later in disease is this followed by loss of vGlut2-positive RGC axon terminals (Crish et al., 2010). It remains to be determined whether changes in synaptic function and neuronal excitability accompanying OHT follow a similar distal-to-proximal progression. If transport defects and synaptic dysfunction are part of the same cascade of OHT-triggered optic nerve defects, we expect that to be the case.

Beyond changes in dendritic transport, other rodent studies have pointed to a shrinkage of neurons in the dLGN and superior colliculus (SC), loss of dLGN neurons, and changes in SC receptive fields (Chen et al., 2015; Wong and Brown, 2012; Zhang et al., 2009). These patterns are largely paralleled in studies from nonhuman primates (Gupta et al., 2007; Ito et al., 2011; Liu et al., 2011; Ly et al., 2011; Shimazawa et al., 2012; Yücel et al., 2000, 2001, 2003). Histology of post-mortem human patient tissue and MRI imaging of human subjects also show similar neuronal atrophy and decreases in LGN size indicating that these effects are a hallmark of the disease rather than a quirk of the experimental system (Chaturvedi et al., 1993; Gupta et al., 2006, 2009).

Different regions of the human and primate LGN are differentially affected in glaucoma, with koniocellular layers experiencing an earlier somatic shrinkage and neuronal loss than magnocellular and parvocellular layers (Gupta et al., 2006; Yücel et al., 2000, 2003) and differential effects on neuronal loss in magnocellular vs. parvocellular layers (Chaturvedi et al., 1993). The mouse dLGN has a simpler arrangement than primate (Kerschensteiner and Guido, 2017). Eye-specific inputs are largely segregated, with axons from contralateral RGCs making up the bulk of the input to the dLGN. Analogous to the primate LGN, different dLGN regions in rodent also receive input from different RGC classes and therefore appear to be responsible for distinct visual information channels. The shell region, which is located on the dorsal surface, receives input principally from On-Off direction-selective RGCs while the core receives input from On and Off sustained and transient RGCs (Kerschensteiner and Guido, 2017).

In this study, we measured the influence of OHT on retinogeniculate synaptic transmission using a reporter line in which ChR2 was used to selectively excite the axons of several classes of melanopsin-expressing retinal ganglion cells (Ecker et al., 2010; Estevez et al., 2012; Schmidt et al., 2014; Sonoda et al., 2018; Stabio et al., 2018). Although the “classical” melanopsin-expressing RGCs (M1 cells) are most famously responsible for non-image-forming vision and do not send axons to the dLGN (Berson et al., 2002; Hattar et al., 2002, 2006), other populations (M4 and M5’s) have less melanopsin, project to the dLGN, and therefore contribute to conscious vision (Ecker et al., 2010; Estevez et al., 2012; Schmidt et al., 2014; Sonoda et al., 2018; Stabio et al., 2018). Since we recorded from the dLGN core, a region that receives substantial input from these neurons (Ecker et al., 2010; Estevez et al., 2012), the results of this study pertain to M4 and M5 RGCs rather than M1 cells. Interestingly, experimental evidence in a variety of animal glaucoma models has suggested that melanopsin-expressing cells are uniquely resistant to optic nerve injury (Cui et al., 2015). It remains to be seen whether OHT has stronger effects on the outputs of RGCs that do not express melanopsin, or whether optic nerve injury is non-selective for RGC type at their dLGN outputs. If different RGCs and their associated visual pathways are affected at different time courses in glaucoma, then designing stimuli to carefully probe those pathways might be a viable approach for early glaucoma diagnosis.

The mechanisms linking OHT to changes in neuronal structure and function in the brain are still unclear. OHT causes an increase in astrocytes and microglia in the dLGN (Shimazawa et al., 2012; Zhang et al., 2009). Microglia are known to mediate dendritic pruning in development and disease (Stephan et al., 2012; Stevens et al., 2007) and so might contribute to dLGN relay neuron dendritic remodeling. NMDA-type glutamate receptors (NMDARs) can contribute to Ca^2+^-dependent excitotoxicity (Lau and Tymianski, 2010) and are present at retinogeniculate synapses (Chen and Regehr, 2000; Lo et al., 2002). NMDAR inhibition by oral dosing of memantine reduced some of the effects of OHT on dendritic complexity in a primate glaucoma model (Yücel et al., 2006). Changes in visual activity triggers changes in Ca^2+^-permeable AMPAR expression at retinogeniculate synapses (Louros et al., 2014) and cp-AMPARs also contribute to excitotoxicity (Wang et al., 2014). It is unclear if this occurs at retinogeniculate synapses during OHT; cp-AMPARs have a higher single-channel conductance (Swanson et al., 1997), which might also lead to an increase in mEPSC amplitude. We did not see any such change in our recordings.

One likely potential mediator of the OHT-triggered changes in retinogeniculate function might be the neurotrophin brain-derived neurotrophic factor (BDNF). Changes in BDNF levels are key for regulating homeostatic plasticity of cortical neurons (Desai et al., 1999; Rutherford et al., 1997, 1998) and BDNF activation of its receptor TrkB are key regulators of long-term synaptic plasticity (Park and Poo, 2013; Poo, 2001; Sasi et al., 2017). While BDNF is found in RGC axons in the dLGN, but not thalamocortical relay (TC) neurons, TrkB is found on both RGC axons and TC neurons (Avwenagha et al., 2006). BDNF levels and transport are altered with age and optic nerve injury including glaucoma, and TrkB activation and transport is also reduced (Avwenagha et al., 2006; Dekeyster et al., 2015; Duprey-Díaz et al., 2002; Gupta et al., 2014; Pease et al., 2000; Quigley et al., 2000). BDNF has also been shown to be important in RGC survival in OHT, as BDNF supplementation can preserve RGCs in rodent ocular hypertension models (Feng et al., 2016; Valiente-Soriano et al., 2015a). The role for BDNF in OHT-triggered changes to retinogeniculate function remains to be tested.

In the DBA/2J mouse, OHT causes defects in optic nerve nodes of ranvier and conveyance of signals to the superior colliculus (Smith et al., 2018). This is accompanied by atrophy of RGC synaptic terminals in the superior colliculus, as measured with 3D scanning block face electron microscopy (Smith et al., 2016). This contrasts with the lack of effect on vGlut2 puncta size measured using light microscopy in our study. Comparison of DBA/2J and microbead-injected mice is challenging, although the difference might also arise from different time points in the disease process or from different effects observed in SC vs dLGN. Smith and colleagues also showed changes in the size and number of presynaptic mitochondria and active zones in the SC. If OHT has similar effects in the dLGN, this might contribute to our observed effects on synaptic function, especially as altered mitochondrial function can lead to changes in presynaptic Ca^2+^ dynamics regulating synaptic vesicle release probability.

Our findings represent a new contribution to the current understanding of OHT-triggered changes in brain function. While somatic atrophy, dendritic remodeling, and neuronal loss all generally lag behind RGC axonal degeneration (Calkins, 2012; Crish et al., 2010; Ito et al., 2011; Shimazawa et al., 2012; Yücel et al., 2001, 2003; Zhang et al., 2009), we have demonstrated that functional changes in the dLGN occur much earlier in disease progression: following IOP elevation, but prior to substantial RGC degeneration. Future work should explore the mechanisms underlying these effects; whether they are triggered by disrupted BDNF signaling and/or altered Ca^2+^ entry and handling and how they relate to the time course of gliosis. Additionally, future work will need to more fully explore the consequences of these changes on different RGC populations and different dLGN subregions.

## Acknowledgements

Scott Nawy for helpful feedback on the manuscript. David Calkins, Wendi Lambert & Melissa Cooper for training in the microbead occlusion method.

## Author contributions

MVH: conceptualization. AB, DG, SF, LR, TG, YZ, JS, MVH: conducting experiments, analyzing data. MVH: Manuscript writing and figure preparation. MVH: supervision and funding.

## References

Abbott, C. J., Choe, T. E., Lusardi, T. A., Burgoyne, C. F., Wang, L., and Fortune, B. (2014). Evaluation of retinal nerve fiber layer thickness and axonal transport 1 and 2 weeks after 8 hours of acute intraocular pressure elevation in rats. Invest. Ophthalmol. Vis. Sci. 55, 674–687. doi:10.1167/iovs.13-12811.

Applebury, M. L., Antoch, M. P., Baxter, L. C., Chun, L. L., Falk, J. D., Farhangfar, F., et al. (2000). The murine cone photoreceptor: a single cone type expresses both S and M opsins with retinal spatial patterning. Neuron 27, 513–523.

Avwenagha, O., Bird, M. M., Lieberman, A. R., Yan, Q., and Campbell, G. (2006). Patterns of expression of brain-derived neurotrophic factor and tyrosine kinase B mRNAs and distribution and ultrastructural localization of their proteins in the visual pathway of the adult rat. Neuroscience 140, 913–928. doi:10.1016/j.neuroscience.2006.02.056.

Berson, D. M., Dunn, F. A., and Takao, M. (2002). Phototransduction by retinal ganglion cells that set the circadian clock. Science 295, 1070–1073. doi:10.1126/science.1067262.

Budisantoso, T., Matsui, K., Kamasawa, N., Fukazawa, Y., and Shigemoto, R. (2012). Mechanisms underlying signal filtering at a multisynapse contact. J. Neurosci. Off. J. Soc. Neurosci. 32, 2357–2376. doi:10.1523/JNEUROSCI.5243-11.2012.

Calkins, D. J. (2012). Critical pathogenic events underlying progression of neurodegeneration in glaucoma. Prog. Retin. Eye Res. 31, 702–719. doi:10.1016/j.preteyeres.2012.07.001.

Calkins, D. J., Lambert, W. S., Formichella, C. R., McLaughlin, W. M., and Sappington, R. M. (2018). The Microbead Occlusion Model of Ocular Hypertension in Mice. Methods Mol. Biol. Clifton NJ 1695, 23–39. doi:10.1007/978-1-4939-7407-8_3.

Chaturvedi, N., Hedley-Whyte, E. T., and Dreyer, E. B. (1993). Lateral geniculate nucleus in glaucoma. Am. J. Ophthalmol. 116, 182–188.

Chen, C., and Regehr, W. G. (2000). Developmental remodeling of the retinogeniculate synapse. Neuron 28, 955–966.

Chen, H., Wei, X., Cho, K.-S., Chen, G., Sappington, R., Calkins, D. J., et al. (2011). Optic neuropathy due to microbead-induced elevated intraocular pressure in the mouse. Invest. Ophthalmol. Vis. Sci. 52, 36–44. doi:10.1167/iovs.09-5115.

Chen, H., Zhao, Y., Liu, M., Feng, L., Puyang, Z., Yi, J., et al. (2015). Progressive degeneration of retinal and superior collicular functions in mice with sustained ocular hypertension. Invest. Ophthalmol. Vis. Sci. 56, 1971–1984. doi:10.1167/iovs.14-15691.

Cooper, M. L., Crish, S. D., Inman, D. M., Horner, P. J., and Calkins, D. J. (2016). Early astrocyte redistribution in the optic nerve precedes axonopathy in the DBA/2J mouse model of glaucoma. Exp. Eye Res. 150, 22–33. doi:10.1016/j.exer.2015.11.016.

Crish, S. D., and Calkins, D. J. (2015). Central visual pathways in glaucoma: evidence for distal mechanisms of neuronal self-repair. J. Neuro-Ophthalmol. Off. J. North Am. Neuro-Ophthalmol. Soc. 35 Suppl 1, S29–37. doi:10.1097/WNO.0000000000000291.

Crish, S. D., Dapper, J. D., MacNamee, S. E., Balaram, P., Sidorova, T. N., Lambert, W. S., et al. (2013). Failure of axonal transport induces a spatially coincident increase in astrocyte BDNF prior to synapse loss in a central target. Neuroscience 229, 55–70. doi:10.1016/j.neuroscience.2012.10.069.

Crish, S. D., Sappington, R. M., Inman, D. M., Horner, P. J., and Calkins, D. J. (2010). Distal axonopathy with structural persistence in glaucomatous neurodegeneration. Proc. Natl. Acad. Sci. U. S. A. 107, 5196–5201. doi:10.1073/pnas.0913141107.

Cui, Q., Ren, C., Sollars, P. J., Pickard, G. E., and So, K.-F. (2015). The injury resistant ability of melanopsin-expressing intrinsically photosensitive retinal ganglion cells. Neuroscience 284, 845–853. doi:10.1016/j.neuroscience.2014.11.002.

Dekeyster, E., Geeraerts, E., Buyens, T., Van den Haute, C., Baekelandt, V., De Groef, L., et al. (2015). Tackling Glaucoma from within the Brain: An Unfortunate Interplay of BDNF and TrkB. PloS One 10, e0142067. doi:10.1371/journal.pone.0142067.

Della Santina, L., Inman, D. M., Lupien, C. B., Horner, P. J., and Wong, R. O.L. (2013). Differential progression of structural and functional alterations in distinct retinal ganglion cell types in a mouse model of glaucoma. J. Neurosci. Off. J. Soc. Neurosci. 33, 17444–17457. doi:10.1523/JNEUROSCI.5461-12.2013.

Della Santina, L., and Ou, Y. (2016). Who’s lost first? Susceptibility of retinal ganglion cell types in experimental glaucoma. Exp. Eye Res. doi:10.1016/j.exer.2016.06.006.

Desai, N. S., Rutherford, L. C., and Turrigiano, G. G. (1999). BDNF regulates the intrinsic excitability of cortical neurons. Learn. Mem. Cold Spring Harb. N 6, 284–291.

Duprey-Díaz, M. V., Soto, I., Blagburn, J. M., and Blanco, R. E. (2002). Changes in brain-derived neurotrophic factor and trkB receptor in the adult Rana pipiens retina and optic tectum after optic nerve injury. J. Comp. Neurol. 454, 456–469. doi:10.1002/cne.10451.

Dzyubenko, E., Rozenberg, A., Hermann, D. M., and Faissner, A. (2016). Colocalization of synapse marker proteins evaluated by STED-microscopy reveals patterns of neuronal synapse distribution in vitro. J. Neurosci. Methods 273, 149–159. doi:10.1016/j.jneumeth.2016.09.001.

Ecker, J. L., Dumitrescu, O. N., Wong, K. Y., Alam, N. M., Chen, S.-K., LeGates, T., et al. (2010). Melanopsin-expressing retinal ganglion-cell photoreceptors: cellular diversity and role in pattern vision. Neuron 67, 49–60. doi:10.1016/j.neuron.2010.05.023.

El-Danaf, R. N., and Huberman, A. D. (2015). Characteristic patterns of dendritic remodeling in early-stage glaucoma: evidence from genetically identified retinal ganglion cell types. J. Neurosci. Off. J. Soc. Neurosci. 35, 2329–2343. doi:10.1523/JNEUROSCI.1419-14.2015.

Estevez, M. E., Fogerson, P. M., Ilardi, M. C., Borghuis, B. G., Chan, E., Weng, S., et al. (2012). Form and function of the M4 cell, an intrinsically photosensitive retinal ganglion cell type contributing to geniculocortical vision. J. Neurosci. Off. J. Soc. Neurosci. 32, 13608–13620. doi:10.1523/JNEUROSCI.1422-12.2012.

Feng, L., Chen, H., Yi, J., Troy, J. B., Zhang, H. F., and Liu, X. (2016). Long-Term Protection of Retinal Ganglion Cells and Visual Function by Brain-Derived Neurotrophic Factor in Mice With Ocular Hypertension. Invest. Ophthalmol. Vis. Sci. 57, 3793–3802. doi:10.1167/iovs.16-19825.

Ferreira, T. A., Blackman, A. V., Oyrer, J., Jayabal, S., Chung, A. J., Watt, A. J., et al. (2014). Neuronal morphometry directly from bitmap images. Nat. Methods 11, 982–984. doi:10.1038/nmeth.3125.

Frankfort, B. J., Khan, A. K., Tse, D. Y., Chung, I., Pang, J.-J., Yang, Z., et al. (2013). Elevated intraocular pressure causes inner retinal dysfunction before cell loss in a mouse model of experimental glaucoma. Invest. Ophthalmol. Vis. Sci. 54, 762–770. doi:10.1167/iovs.12-10581.

Fujiyama, F., Hioki, H., Tomioka, R., Taki, K., Tamamaki, N., Nomura, S., et al. (2003). Changes of immunocytochemical localization of vesicular glutamate transporters in the rat visual system after the retinofugal denervation. J. Comp. Neurol. 465, 234–249. doi:10.1002/cne.10848.

Gazzard, G., Foster, P. J., Devereux, J. G., Oen, F., Chew, P., Khaw, P. T., et al. (2003). Intraocular pressure and visual field loss in primary angle closure and primary open angle glaucomas. Br. J. Ophthalmol. 87, 720–725.

Gupta, N., Ang, L.-C., Noël de Tilly, L., Bidaisee, L., and Yücel, Y. H. (2006). Human glaucoma and neural degeneration in intracranial optic nerve, lateral geniculate nucleus, and visual cortex. Br. J. Ophthalmol. 90, 674–678. doi:10.1136/bjo.2005.086769.

Gupta, N., Greenberg, G., de Tilly, L. N., Gray, B., Polemidiotis, M., and Yücel, Y. H. (2009). Atrophy of the lateral geniculate nucleus in human glaucoma detected by magnetic resonance imaging. Br. J. Ophthalmol. 93, 56–60. doi:10.1136/bjo.2008.138172.

Gupta, N., Ly, T., Zhang, Q., Kaufman, P. L., Weinreb, R. N., and Yücel, Y. H. (2007). Chronic ocular hypertension induces dendrite pathology in the lateral geniculate nucleus of the brain. Exp. Eye Res. 84, 176–184. doi:10.1016/j.exer.2006.09.013.

Gupta, V., You, Y., Li, J., Gupta, V., Golzan, M., Klistorner, A., et al. (2014). BDNF impairment is associated with age-related changes in the inner retina and exacerbates experimental glaucoma. Biochim. Biophys. Acta 1842, 1567–1578. doi:10.1016/j.bbadis.2014.05.026.

Hattar, S., Kumar, M., Park, A., Tong, P., Tung, J., Yau, K.-W., et al. (2006). Central projections of melanopsin-expressing retinal ganglion cells in the mouse. J. Comp. Neurol. 497, 326–349. doi:10.1002/cne.20970.

Hattar, S., Liao, H. W., Takao, M., Berson, D. M., and Yau, K. W. (2002). Melanopsin-containing retinal ganglion cells: architecture, projections, and intrinsic photosensitivity. Science 295, 1065–1070. doi:10.1126/science.1069609.

Ito, Y., Shimazawa, M., Inokuchi, Y., Yamanaka, H., Tsuruma, K., Imamura, K., et al. (2011). Involvement of endoplasmic reticulum stress on neuronal cell death in the lateral geniculate nucleus in the monkey glaucoma model. Eur. J. Neurosci. 33, 843–855. doi:10.1111/j.1460-9568.2010.07578.x.

Kerschensteiner, D., and Guido, W. (2017). Organization of the dorsal lateral geniculate nucleus in the mouse. Vis. Neurosci. 34, E008. doi:10.1017/S0952523817000062.

Krahe, T. E., El-Danaf, R. N., Dilger, E. K., Henderson, S. C., and Guido, W. (2011). Morphologically distinct classes of relay cells exhibit regional preferences in the dorsal lateral geniculate nucleus of the mouse. J. Neurosci. Off. J. Soc. Neurosci. 31, 17437–17448. doi:10.1523/JNEUROSCI.4370-11.2011.

Land, P. W., Kyonka, E., and Shamalla-Hannah, L. (2004). Vesicular glutamate transporters in the lateral geniculate nucleus: expression of VGLUT2 by retinal terminals. Brain Res. 996, 251–254.

Lau, A., and Tymianski, M. (2010). Glutamate receptors, neurotoxicity and neurodegeneration. Pflugers Arch. 460, 525–542. doi:10.1007/s00424-010-0809-1.

Lin, D. J., Kang, E., and Chen, C. (2014). Changes in input strength and number are driven by distinct mechanisms at the retinogeniculate synapse. J. Neurophysiol. 112, 942–950. doi:10.1152/jn.00175.2014.

Litvina, E. Y., and Chen, C. (2017). Functional Convergence at the Retinogeniculate Synapse. Neuron 96, 330–338.e5. doi:10.1016/j.neuron.2017.09.037.

Liu, H.-H., Zhang, L., Shi, M., Chen, L., and Flanagan, J. G. (2017). Comparison of laser and circumlimbal suture induced elevation of intraocular pressure in albino CD-1 mice. PloS One 12, e0189094. doi:10.1371/journal.pone.0189094.

Liu, M., Duggan, J., Salt, T. E., and Cordeiro, M. F. (2011). Dendritic changes in visual pathways in glaucoma and other neurodegenerative conditions. Exp. Eye Res. 92, 244–250. doi:10.1016/j.exer.2011.01.014.

Liu, M., Guo, L., Salt, T. E., and Cordeiro, M. F. (2014). Dendritic changes in rat visual pathway associated with experimental ocular hypertension. Curr. Eye Res. 39, 953–963. doi:10.3109/02713683.2014.884594.

Lo, F.-S., Ziburkus, J., and Guido, W. (2002). Synaptic mechanisms regulating the activation of a Ca(2+)-mediated plateau potential in developing relay cells of the LGN. J. Neurophysiol. 87, 1175–1185. doi:10.1152/jn.00715.1999.

Longair, M. H., Baker, D. A., and Armstrong, J. D. (2011). Simple Neurite Tracer: open source software for reconstruction, visualization and analysis of neuronal processes. Bioinforma. Oxf. Engl. 27, 2453–2454. doi:10.1093/bioinformatics/btr390.

Louros, S. R., Hooks, B. M., Litvina, L., Carvalho, A. L., and Chen, C. (2014). A role for stargazin in experience-dependent plasticity. Cell Rep. 7, 1614–1625. doi:10.1016/j.celrep.2014.04.054.

Ly, T., Gupta, N., Weinreb, R. N., Kaufman, P. L., and Yücel, Y. H. (2011). Dendrite plasticity in the lateral geniculate nucleus in primate glaucoma. Vision Res. 51, 243–250. doi:10.1016/j.visres.2010.08.003.

Madisen, L., Mao, T., Koch, H., Zhuo, J., Berenyi, A., Fujisawa, S., et al. (2012). A toolbox of Cre-dependent optogenetic transgenic mice for light-induced activation and silencing. Nat. Neurosci. 15, 793–802. doi:10.1038/nn.3078.

Mao, L. K., Stewart, W. C., and Shields, M. B. (1991). Correlation between intraocular pressure control and progressive glaucomatous damage in primary open-angle glaucoma. Am. J. Ophthalmol. 111, 51–55.

Martin, K. R. G., Quigley, H. A., Valenta, D., Kielczewski, J., and Pease, M. E. (2006). Optic nerve dynein motor protein distribution changes with intraocular pressure elevation in a rat model of glaucoma. Exp. Eye Res. 83, 255–262. doi:10.1016/j.exer.2005.11.025.

Miglior, S., and Bertuzzi, F. (2013). Relationship between intraocular pressure and glaucoma onset and progression. Curr. Opin. Pharmacol. 13, 32–35. doi:10.1016/j.coph.2012.09.014.

Morgan, J. L., Berger, D. R., Wetzel, A. W., and Lichtman, J. W. (2016). The Fuzzy Logic of Network Connectivity in Mouse Visual Thalamus. Cell 165, 192–206. doi:10.1016/j.cell.2016.02.033.

Novak, A., Guo, C., Yang, W., Nagy, A., and Lobe, C. G. (2000). Z/EG, a double reporter mouse line that expresses enhanced green fluorescent protein upon Cre-mediated excision. Genes. N. Y. N 2000 28, 147–155.

Ou, Y., Jo, R. E., Ullian, E. M., Wong, R. O.L., and Della Santina, L. (2016). Selective Vulnerability of Specific Retinal Ganglion Cell Types and Synapses after Transient Ocular Hypertension. J. Neurosci. Off. J. Soc. Neurosci. 36, 9240–9252. doi:10.1523/JNEUROSCI.0940-16.2016.

Pang, J.-J., Frankfort, B. J., Gross, R. L., and Wu, S. M. (2015). Elevated intraocular pressure decreases response sensitivity of inner retinal neurons in experimental glaucoma mice. Proc. Natl. Acad. Sci. U. S. A. 112, 2593–2598. doi:10.1073/pnas.1419921112.

Park, H., and Poo, M. (2013). Neurotrophin regulation of neural circuit development and function. Nat. Rev. Neurosci. 14, 7–23. doi:10.1038/nrn3379.

Park, H.-Y. L., Kim, J. H., and Park, C. K. (2014). Alterations of the synapse of the inner retinal layers after chronic intraocular pressure elevation in glaucoma animal model. Mol. Brain 7, 53. doi:10.1186/s13041-014-0053-2.

Pease, M. E., McKinnon, S. J., Quigley, H. A., Kerrigan-Baumrind, L. A., and Zack, D. J. (2000). Obstructed axonal transport of BDNF and its receptor TrkB in experimental glaucoma. Invest. Ophthalmol. Vis. Sci. 41, 764–774.

Poo, M. M. (2001). Neurotrophins as synaptic modulators. Nat. Rev. Neurosci. 2, 24–32. doi:10.1038/35049004.

Quattrochi, L. E., Stabio, M. E., Kim, I., Ilardi, M. C., Michelle Fogerson, P., Leyrer, M. L., et al. (2019). The M6 cell: A small-field bistratified photosensitive retinal ganglion cell. J. Comp. Neurol. 527, 297–311. doi:10.1002/cne.24556.

Quigley, H. A., McKinnon, S. J., Zack, D. J., Pease, M. E., Kerrigan-Baumrind, L. A., Kerrigan, D. F., et al. (2000). Retrograde axonal transport of BDNF in retinal ganglion cells is blocked by acute IOP elevation in rats. Invest. Ophthalmol. Vis. Sci. 41, 3460–3466.

Rafols, J. A., and Valverde, F. (1973). The structure of the dorsal lateral geniculate nucleus in the mouse. A Golgi and electron microscopic study. J. Comp. Neurol. 150, 303–332. doi:10.1002/cne.901500305.

Risner, M. L., Pasini, S., Cooper, M. L., Lambert, W. S., and Calkins, D. J. (2018). Axogenic mechanism enhances retinal ganglion cell excitability during early progression in glaucoma. Proc. Natl. Acad. Sci. U. S. A. 115, E2393–E2402. doi:10.1073/pnas.1714888115.

Rodriguez, A. R., de Sevilla Müller, L. P., and Brecha, N. C. (2014). The RNA binding protein RBPMS is a selective marker of ganglion cells in the mammalian retina. J. Comp. Neurol. 522, 1411–1443. doi:10.1002/cne.23521.

Rutherford, L. C., DeWan, A., Lauer, H. M., and Turrigiano, G. G. (1997). Brain-derived neurotrophic factor mediates the activity-dependent regulation of inhibition in neocortical cultures. J. Neurosci. Off. J. Soc. Neurosci. 17, 4527–4535.

Rutherford, L. C., Nelson, S. B., and Turrigiano, G. G. (1998). BDNF has opposite effects on the quantal amplitude of pyramidal neuron and interneuron excitatory synapses. Neuron 21, 521–530.

Sakaba, T., Schneggenburger, R., and Neher, E. (2002). Estimation of quantal parameters at the calyx of Held synapse. Neurosci. Res. 44, 343–356.

Salvadores, N., Sanhueza, M., Manque, P., and Court, F. A. (2017). Axonal Degeneration during Aging and Its Functional Role in Neurodegenerative Disorders. Front. Neurosci. 11, 451. doi:10.3389/fnins.2017.00451.

Sappington, R. M., Carlson, B. J., Crish, S. D., and Calkins, D. J. (2010). The microbead occlusion model: a paradigm for induced ocular hypertension in rats and mice. Invest. Ophthalmol. Vis. Sci. 51, 207–216. doi:10.1167/iovs.09-3947.

Sasi, M., Vignoli, B., Canossa, M., and Blum, R. (2017). Neurobiology of local and intercellular BDNF signaling. Pflugers Arch. 469, 593–610. doi:10.1007/s00424-017-1964-4.

Schmidt, T. M., Alam, N. M., Chen, S., Kofuji, P., Li, W., Prusky, G. T., et al. (2014). A role for melanopsin in alpha retinal ganglion cells and contrast detection. Neuron 82, 781–788. doi:10.1016/j.neuron.2014.03.022.

Schmidt, T. M., and Kofuji, P. (2011). An isolated retinal preparation to record light response from genetically labeled retinal ganglion cells. J. Vis. Exp. JoVE. doi:10.3791/2367.

Shimazawa, M., Ito, Y., Inokuchi, Y., Yamanaka, H., Nakanishi, T., Hayashi, T., et al. (2012). An alteration in the lateral geniculate nucleus of experimental glaucoma monkeys: in vivo positron emission tomography imaging of glial activation. PloS One 7, e30526. doi:10.1371/journal.pone.0030526.

Smith, M. A., Xia, C. Z., Dengler-Crish, C. M., Fening, K. M., Inman, D. M., Schofield, B. R., et al. (2016). Persistence of intact retinal ganglion cell terminals after axonal transport loss in the DBA/2J mouse model of glaucoma. J. Comp. Neurol. 524, 3503–3517. doi:10.1002/cne.24012.

Smith, M.A., Plyler, E.S., Dengler-Crish, C.M., Meier, J., Crish, S.D. (2018). Nodes of Ranvier in glaucoma. Neuroscience. 300, 104–118. doi:10.1016/j.neuroscience.2018.08.016.

Sondereker, K. B., Stabio, M. E., Jamil, J. R., Tarchick, M. J., and Renna, J. M. (2018). Where You Cut Matters: A Dissection and Analysis Guide for the Spatial Orientation of the Mouse Retina from Ocular Landmarks. J. Vis. Exp. doi:10.3791/57861.

Sonoda, T., Lee, S. K., Birnbaumer, L., and Schmidt, T. M. (2018). Melanopsin Phototransduction Is Repurposed by ipRGC Subtypes to Shape the Function of Distinct Visual Circuits. Neuron. doi:10.1016/j.neuron.2018.06.032.

Stabio, M. E., Sabbah, S., Quattrochi, L. E., Ilardi, M. C., Fogerson, P. M., Leyrer, M. L., et al. (2018). The M5 Cell: A Color-Opponent Intrinsically Photosensitive Retinal Ganglion Cell. Neuron 97, 150–163.e4. doi:10.1016/j.neuron.2017.11.030.

Stephan, A. H., Barres, B. A., and Stevens, B. (2012). The complement system: an unexpected role in synaptic pruning during development and disease. Annu. Rev. Neurosci. 35, 369–389. doi:10.1146/annurev-neuro-061010-113810.

Stevens, B., Allen, N. J., Vazquez, L. E., Howell, G. R., Christopherson, K. S., Nouri, N., et al. (2007). The classical complement cascade mediates CNS synapse elimination. Cell 131, 1164–1178. doi:10.1016/j.cell.2007.10.036.

Swanson, G. T., Kamboj, S. K., and Cull-Candy, S. G. (1997). Single-channel properties of recombinant AMPA receptors depend on RNA editing, splice variation, and subunit composition. J. Neurosci. Off. J. Soc. Neurosci. 17, 58–69.

Ting, J. T., Daigle, T. L., Chen, Q., and Feng, G. (2014). Acute brain slice methods for adult and aging animals: application of targeted patch clamp analysis and optogenetics. Methods Mol. Biol. Clifton NJ 1183, 221–242. doi:10.1007/978-1-4939-1096-0_14.

Ting, J. T., Lee, B. R., Chong, P., Soler-Llavina, G., Cobbs, C., Koch, C., et al. (2018). Preparation of Acute Brain Slices Using an Optimized N-Methyl-D-glucamine Protective Recovery Method. J. Vis. Exp. JoVE. doi:10.3791/53825.

Turner, J.P., Salt, T.E. (1998). Characterization of sensory and corticothalamic excitatory inputs to rat thalamocortical neurones in vitro. J. Physiol. 510, 829–843. doi:10.1111/j.1469-7793.1998.829bj.x.

Valiente-Soriano, F. J., Nadal-Nicolás, F. M., Salinas-Navarro, M., Jiménez-López, M., Bernal-Garro, J. M., Villegas-Pérez, M. P., et al. (2015a). BDNF Rescues RGCs But Not Intrinsically Photosensitive RGCs in Ocular Hypertensive Albino Rat Retinas. Invest. Ophthalmol. Vis. Sci. 56, 1924–1936. doi:10.1167/iovs.15-16454.

Valiente-Soriano, F. J., Salinas-Navarro, M., Jiménez-López, M., Alarcón-Martínez, L., Ortín-Martínez, A., Bernal-Garro, J. M., et al. (2015b). Effects of ocular hypertension in the visual system of pigmented mice. PloS One 10, e0121134. doi:10.1371/journal.pone.0121134.

Wang, A. L., Carroll, R. C., and Nawy, S. (2014). Down-regulation of the RNA editing enzyme ADAR2 contributes to RGC death in a mouse model of glaucoma. PloS One 9, e91288. doi:10.1371/journal.pone.0091288.

Weinreb, R. N., Aung, T., and Medeiros, F. A. (2014). The pathophysiology and treatment of glaucoma: a review. JAMA 311, 1901–1911. doi:10.1001/jama.2014.3192.

Wong, A. A., and Brown, R. E. (2012). A neurobehavioral analysis of the prevention of visual impairment in the DBA/2J mouse model of glaucoma. Invest. Ophthalmol. Vis. Sci. 53, 5956–5966. doi:10.1167/iovs.12-10020.

Yucel, Y. H., and Gupta, N. (2015). A framework to explore the visual brain in glaucoma with lessons from models and man. Exp. Eye Res. 141, 171–178. doi:10.1016/j.exer.2015.07.004.

Yücel, Y. H., Gupta, N., Zhang, Q., Mizisin, A. P., Kalichman, M. W., and Weinreb, R. N. (2006). Memantine protects neurons from shrinkage in the lateral geniculate nucleus in experimental glaucoma. Arch. Ophthalmol. Chic. Ill 1960 124, 217–225. doi:10.1001/archopht.124.2.217.

Yücel, Y. H., Zhang, Q., Gupta, N., Kaufman, P. L., and Weinreb, R. N. (2000). Loss of neurons in magnocellular and parvocellular layers of the lateral geniculate nucleus in glaucoma. Arch. Ophthalmol. Chic. Ill 1960 118, 378–384.

Yücel, Y. H., Zhang, Q., Weinreb, R. N., Kaufman, P. L., and Gupta, N. (2001). Atrophy of relay neurons in magno- and parvocellular layers in the lateral geniculate nucleus in experimental glaucoma. Invest. Ophthalmol. Vis. Sci. 42, 3216–3222.

Yücel, Y. H., Zhang, Q., Weinreb, R. N., Kaufman, P. L., and Gupta, N. (2003). Effects of retinal ganglion cell loss on magno-, parvo-, koniocellular pathways in the lateral geniculate nucleus and visual cortex in glaucoma. Prog. Retin. Eye Res. 22, 465–481.

Zhang, S., Wang, H., Lu, Q., Qing, G., Wang, N., Wang, Y., et al. (2009). Detection of early neuron degeneration and accompanying glial responses in the visual pathway in a rat model of acute intraocular hypertension. Brain Res. 1303, 131–143. doi:10.1016/j.brainres.2009.09.029.

Zhuang, X., Sun, W., and Xu-Friedman, M. A. (2017). Changes in Properties of Auditory Nerve Synapses following Conductive Hearing Loss. J. Neurosci. Off. J. Soc. Neurosci. 37, 323–332. doi:10.1523/JNEUROSCI.0523-16.2017.

